# Confirmation of interpersonal expectations is intrinsically rewarding

**DOI:** 10.1101/2020.07.19.210757

**Authors:** Niv Reggev, Anoushka Chowdhary, Jason P. Mitchell

## Abstract

People want to interact successfully with other individuals, and they invest significant efforts in attempting to do so. Decades of research have demonstrated that to simplify the dauntingly complex task of interpersonal communication, perceivers use stereotypes and other sources of prior knowledge to predict the responses of individuals in their environment. Here, we show that these top-down expectations can also shape the subjective value of expectation-consistent and expectation-violating targets. Specifically, in two neuroimaging experiments (*n* = 58), we observed increased activation in brain regions associated with reward processing—including the nucleus accumbens—when perceivers observed information consistent with their social expectations. In two additional behavioral experiments (*n* = 704), we observed that perceivers were willing to forgo money to encounter an expectation-consistent target and avoid an expectation-violating target. Together, these findings suggest that perceivers value having their social expectations confirmed, much like food or monetary rewards.

---

People dedicate a substantial portion of their time to interacting with other individuals. When we succeed in doing so, we feel better physically and psychologically (Baumeister and Leary, 1995; Tay *et al.*, 2013; Cacioppo *et al.*, 2014; Matthews and Tye, 2019). At the same time, other people also present one of the most complicated challenges we have to face. Understanding another person involves inferring hidden states based on fragmentary sensory, verbal and visceral cues, each of which conveys only a small amount of information. To simplify the highly demanding challenge of social cognition, perceivers use top-down predictions (e.g., stereotypes) that help make sense of others in a rapid fashion (Freeman and Johnson, 2016; Otten *et al.*, 2017; Tamir and Thornton, 2018; Hutchinson and Barrett, 2019). Perceivers can thus seamlessly interact with their environment while refraining from the effortful construction of elaborative representations for each individual they encounter (Fiske and Neuberg, 1990; Macrae and Bodenhausen, 2000). For example, on her first day of school, a freshman might assume that her female peers may be interested in conversing about shopping. Utilizing these predictions, the freshman could effortlessly engage in spontaneous conversations with her new female peers, potentially facilitating her social bonds.

However, although they often facilitate interpersonal interaction, social predictions also impose a cost on perceivers. Individuals interpret ambiguous information in accordance with their expectations to confirm existing biases (Darley and Gross, 1983). Moreover, across multiple contexts individuals tend to adhere to predictions they have previously formed and fail to modify them even in the face of contradictory evidence (Hamilton and Sherman, 1996; Gregg *et al.*, 2006; Roese and Sherman, 2007; Wyer, 2010; Dunsmoor *et al.*, 2016). In the person perception domain, when perceivers first learn that someone is ‘intelligent’ and then subsequently learn that he is also ‘envious’, they form a favorably-skewed impression of that person; on the other hand, when perceivers learn about these same two character traits in reverse order, they form an unfavorable impression of the target (Asch, 1946; Sullivan, 2019). In the domain of stereotypes about social groups, perceivers typically do not update self-reported or implicitly measured preferences for a social group once formed (Gregg *et al.*, 2006; Roese and Sherman, 2007). Together, such findings provide further evidence that people prefer to have their social predictions confirmed across multiple domains. To date, research has not been able to identify the sources that support the persistence of these initial predictions about other individuals. Here, we integrate insights from social psychology and neuroscience to explore the idea that perceivers prefer to ‘stick’ with their initial predictions because they attribute subjective value to the confirmation of these predictions. That is, we posit that targets who confirm our expectations about them (such as stereotype-consistent targets) will trigger a reward-like response similar to food, sex, or chocolate.

Several lines of research already hint at such an effect. Perceivers generally like individuals who conform to expectations more than individuals who violate them (Eagly and Karau, 2002; Phelan and Rudman, 2010; Stern *et al.*, 2015); for example, observers typically prefer female teachers to male teachers, but like male leaders better than their equally competent female peers (Rudman *et al.*, 2012; Moss-Racusin and Johnson, 2016). Similarly, participants express greater trust in targets that fit gender-based predictions (Olszanowski *et al.*, 2018; Stern and Rule, 2018). Moreover, perceivers demonstrate similar effects for emotion-based expectations, regardless of the valence of the emotion (Chanes *et al.*, 2018). Several theorists have suggested that perceivers may gradually develop a habitual hedonic response for targets conforming to normative expectations (Jost and Hunyady, 2003; Berridge, 2012; Huebner, 2016; Theriault *et al.*, 2020). These suggestions dovetail with the well-documented aversive reactions people experience when confronted with violations of predictions and the uncertainty associated with such violations (Festinger, 1957; Roese and Sherman, 2007; Gawronski, 2012; FeldmanHall and Shenhav, 2019; Theriault *et al.*, 2020). In a similar vein, when perceivers attempt to form an impression about an expectation-violating social target, they effortfully process the information (Fiske and Neuberg, 1990) and show enhanced activity in several brain regions, including the dorsomedial prefrontal cortex (Cloutier *et al.*, 2011; Ames and Fiske, 2013; Mende-Siedlecki *et al.*, 2013). Put together, these positive and aversive responses motivate perceivers to seek expectation-consistent information.

In spite of ample evidence for perceivers’ motivation to confirm their social expectations, scholars are still debating the mechanisms supporting the persistence of this motivation. Here we suggest that neural activity can offer a novel insight on this topic. In recent years, scholars have identified the involuntary effects of motivation and expectation in several neural systems, most notably in the mesolimbic dopaminergic system (Kohli *et al.*, 2018). Animal models suggest that midbrain dopaminergic activity signals one’s internal desire to obtain a goal (Berridge, 2012). In humans, the motivation to experience positive effects manifests specifically in midbrain and striatal responses to better- or worse-than-expected information (Sharot *et al.*, 2012; Lefebvre *et al.*, 2017; Charpentier *et al.*, 2018). Likewise, participants expecting a painful stimulus demonstrate increased striatal activity while experiencing pain, compared with participants who do not expect to feel pain (Jepma *et al.*, 2018; Schwarz *et al.*, 2019; for a related effect of negative stigma, see Welborn *et al.*, 2020). Finally, a recent meta-analysis reported that when perceivers agreed with expected group opinions, they demonstrated robust striatal activity, as compared with times in which they deviated from the group consensus (Wu *et al.*, 2016). These studies suggest that insofar as perceivers hold a motivation to experience a certain event, the striatum responds to events that align with that motivation.

Notably, the involvement of the striatum hints at a potential mechanism driving the effects of expectation and motivation. Researchers repeatedly identify striatal activity, and most prominently activity in its ventral portion, in anticipation and receipt of various types of reward (Schultz, 2000; Hare *et al.*, 2008; Haber and Knutson, 2010). For example, the nucleus accumbens (NAcc), located at the ventral-rostral tip of the striatum, responds both to primary rewards (e.g., food or erotic) and secondary rewards (e.g., money or positive feedback) (Peters and Büchel, 2010; Bartra *et al.*, 2013; Sescousse *et al.*, 2013). The NAcc also responds to social experiences, such as engagement with attractive or smiling faces, prosocial actions, or placing one’s trust in peers (Harris and Fiske, 2010; Lin *et al.*, 2012; Hackel *et al.*, 2015; Krosch and Amodio, 2019). As increased striatal responses are often associated with rewards, such a response to events aligning with perceiver’s motivation might indicate that these events are rewarding as well.

Together, these studies suggest that consistency with stereotypes and other forms of interpersonal predictions is intrinsically rewarding. To test this hypothesis, we first measured NAcc activity in response to information consistent or inconsistent with social expectations. Using functional magnetic resonance imaging (fMRI), we scanned participants while they judged target individuals on characteristics that either conformed to or violated interpersonal expectations. As the NAcc is consistently involved in rewarding experiences, activity in this region can serve as a marker of a neural reward response. If perceivers value having their social expectations confirmed, we should observe increased NAcc activity for trials in which targets were associated with characteristics that confirm expectations compared to trials in which targets violate them.

In addition, we assessed whether perceivers actively prefer expectancy-confirming social information by creating experimental situations in which participants could trade money for the chance to view targets with expectation-consistent characteristics. To do so, we relied on a modified version of a “pay-per-view” task, previously used with human and non-human primates, to measure the monetary value associated with expectancy-confirming and expectancy-violating stimuli (Deaner *et al.*, 2005; Tamir and Mitchell, 2012). If perceivers experience expectancy-confirming information as intrinsically more valuable, we expected participants to forgo money to rate individuals associated with expectancy-consistent rather than expectancy-violating information.

Interestingly, social expectations can take multiple forms. For example, stereotypes relate specific attributes to social groups regardless of personal knowledge about group members. Conversely, we can construct detailed individuated expectations about familiar individuals, such as personally familiar others or famous people (Fiske and Neuberg, 1990; Hamilton and Sherman, 1996). Accordingly, we also probed whether predictions from these two distinct sources evoked qualitatively different reward responses. Together, the results of four studies support the hypothesis that perceivers are motivated to reaffirm their interpersonal forecasts, in part because they experience the confirmation of social expectations as a powerful form of subjective reward.

## Results

### Study 1: Neural response to stereotype confirmation vs. violation

In Study 1 we observed greater NAcc activity when participants saw targets associated with gender-stereotype-consistent information than when they saw targets associated with stereotype-violating information. We scanned participants (*n* = 28; here and in subsequent studies, the sample size reported includes only participants who did not fail pre-defined exclusion criteria; see Methods section) while they formed impressions about targets that varied in the degree to which they confirmed gender stereotypes. Gender stereotypes included various characteristics typically associated with men (e.g., “emotionally closed” or “CEO of a big company”) or women (e.g., “loves children” or “an admired preschool teacher”). In each of 204 trials, participants first read a short description for 1.5 seconds and then saw the face of a target man or woman (see Fig S1A). Participants rated how likely the target was to have the presented characteristic (see SI results and Table S1 for behavioral results). We conducted three parallel analyses to examine whether the neural region most associated with reward—namely, the NAcc—was more engaged when the target was associated with stereotype-derived expectations compared to when the target violated them. First, a whole-brain random-effects contrast identified regions that were more active for *stereotype-consistent > stereotype-violating* trials (*p* < 0.05, corrected; see Table S2 for full results). This analysis indicated a significantly greater response in the NAcc when the presented target matched the stereotypical expectation set by the preceding statement than when the target violated that expectation (Fig. 1A). Parallel results were obtained when we modeled expectation consistency as a continuous rather than dichotomized predictor (see Table S2 and SI Materials and Methods for full details).

**Figure 1.**
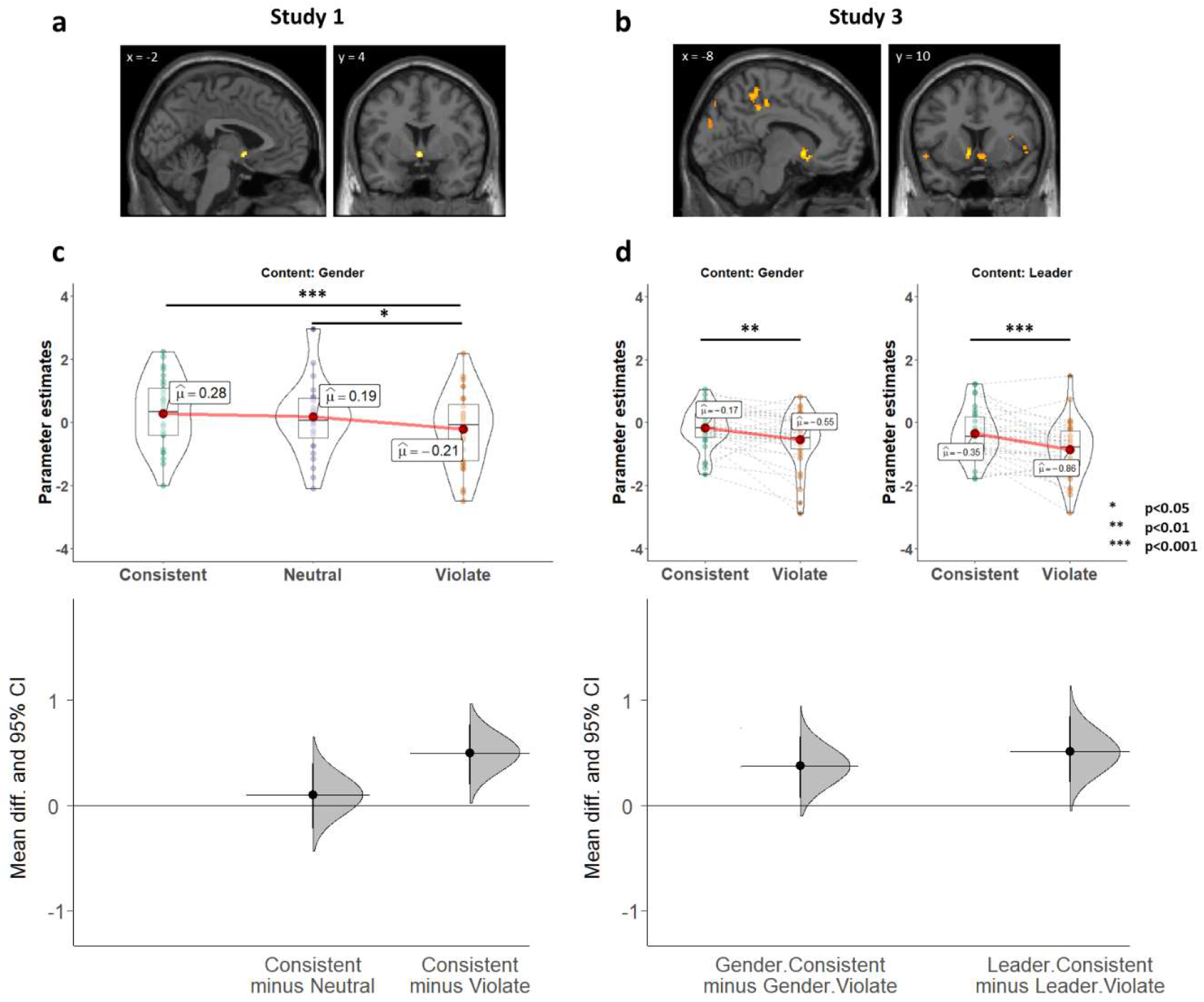
Neural responses associated with the confirmation of expectations about other individuals. (a) Whole-brain random-effects contrasts comparing *expectation-consistent > expectation-violating* trials revealed activity in the nucleus accumbens (NAcc) in (A) Study 1 and (B) Study 3. (C) We independently defined a region of interest in the NAcc using a comprehensive meta-analysis (MNI coordinates: −6, 10, −6; 10, 12, −6). Analysis of parameter estimates in this region confirmed that the bilateral NAcc showed a stronger response during consistent than during violating trials in Study 1 and (D) Study 3. Upper panel: Across all figures, individual dots represent parameter estimates for individual participants. Each figure also visualizes the mean of each condition (as a red dot), the median (solid horizontal line), and first and third quartiles (boxplot). Lower panel: effect size (the mean difference between respective conditions, indicated by the black circles), the bootstrapped 95% confidence intervals (illustrated by the vertical lines), and the resampled distribution of the effect size given the observed data, indicated by the curve (see Methods section).

Second, to confirm that this region overlapped with those responsive to rewards, we independently defined a neural regions of interest (ROI) based on spheres around peak voxels identified in a comprehensive meta-analysis (Bartra *et al.*, 2013). In this independently defined region consistency with stereotypes resulted in significantly greater activity compared to their violation [one-sided test: *t*_(54)_ = 3.53, *p* = 0.0004, *Hedges’s g* = 0.37 [95% confidence intervals: 0.1–0.64], Fig. 1B]. Third, we corroborated this finding by defining ROIs from a task in which participants received monetary rewards based on their performance [the Monetary Incentive Delay (MID) task; see Materials and Methods] (Knutson *et al.*, 2000). This analysis yielded similar results [*t*_(54)_ = 2.87, *p* = 0.0029, *g* = 0.30 [0.03–0.59]]. Together, these patterns suggest that seeing a person associated with a stereotypical characteristic triggers activation in the very same region that responds to primary and secondary reinforcers, highlighting the intrinsic value of stereotype confirmation.

Additionally, to investigate whether this NAcc activation is limited to stereotype-derived expectations, we included a third type of statements in our study: stereotype-neutral statements (e.g., “drinks coffee every morning”). Interestingly, we found that the overall NAcc response to targets associated with stereotype-neutral information was higher than the response to stereotype-violating information and not different from stereotype-consistent information [Fig. 1C; Bonferroni corrected comparisons: neutral versus violating: Meta-analysis ROIs: t_(54)_ = 2.83, p = 0.0195, g = 0.29 [0.09–0.51]; MID ROIs: t_(54)_ = 2.38, p = 0.062, g = 0.25 [0.04–0.47]; neutral versus consistent: Meta-analysis ROIs: t_(54)_ = 0.7, p = 0.76, g = 0.07 [−0.19–0.34]; MID ROIs: t_(54)_ = 0.49, p = 0.63, g = 0.05 [−0.18–0.28]]. At first blush, this finding suggests that NAcc activation was modulated only by stereotype-inconsistent information, not that it was especially driven by stereotype-consistent information which, on average, did not differ from stereotype-neutral trials. However, additional analyses belie this interpretation. On each trial, participants rated the likelihood that the target could be described by the accompanying characteristic (e.g., “enjoys drinking coffee in the morning”). Importantly, the NAcc was activated during stereotype-neutral trials only when participants endorsed those characteristics as descriptive of the target and not when they rejected its applicability to a target (Meta-analysis ROIs: F_(1,76.18)_ = 15.67, p = 0.0002; η_p_^2^ = 0.17 [0.06, 0.29]; MID ROIs: F_(1,76.5)_ = 12.66, p = 0.0006; η_p_^2^ = 0.14 [0.04, 0.26]). In contrast, the preferential NAcc activation observed for stereotype-consistent trials was unaffected by participants’ likelihood ratings (interaction between response and condition: Meta-analysis ROIs: F_(1.77,47.68)_ = 5.4, p = 0.01, η_p_^2^ = 0.17 [0.02, 0.3]; MID ROIs: F_(1.87,50.4)_ = 4.01, p = 0.03, η_p_^2^ = 0.13 [0.01, 0.26]; see Fig. 2. Stereotype-consistent trials: meta-analysis ROIs: F_(1,76.18)_ = 0.2, p > 0.5; η_p_^2^ = 0.003 [0, 0.05]; MID ROIs: F_(1,76.5)_ = 0.63, p > 0.5; η_p_^2^ = 0.005 [0, 0.06]). Together, these data points suggest that NAcc is activated specifically when one’s expectations are confirmed, regardless of whether those expectations derive from culturally-salient stereotypes or from more personal and idiosyncratic sources. Finally, as for stereotype-neutral trials, stereotype-violating trials (e.g., a man who “enjoys shopping for shoes”) were associated with greater NAcc activation when participants endorsed the likelihood of such characteristics (Meta-analysis ROIs: F_(1,76.18)_ = 17.84, p = 0.0001; η_p_^2^ = 0.19 [0.07, 0.31]; MID ROIs: F_(1,76.5)_ = 15.76, p = 0.0002; η_p_^2^ = 0.17 [0.06, 0.29]).

**Figure 2.**
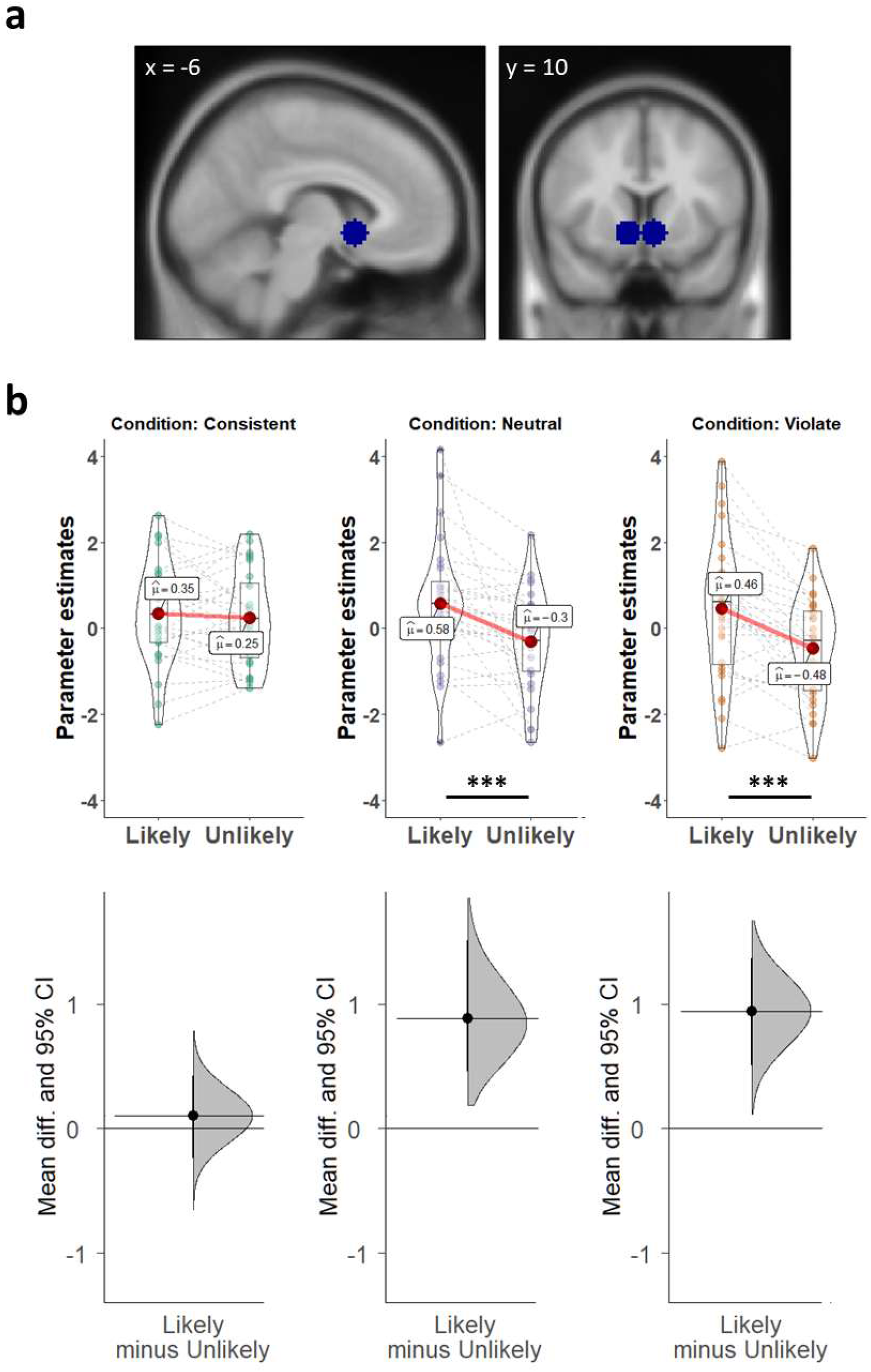
The effect of behavioral ratings of trials on neural responses. In each trial, participants indicated whether they thought the presented target was likely or unlikely to be associated with the presented statement. (A) The independently defined regions of interest in the NAcc. (B) We observed a significant interaction in the independently defined NAcc in Study 1: Participants’ ratings modulated the neural response only for stereotype-neutral and stereotype-violating targets, suggesting that stereotype-confirming targets are involuntarily rewarding.

### Study 2: The monetary value of stereotype confirmation

Although activation of the NAcc often reflects the presence of rewarding stimuli (Bhanji and Delgado, 2014), this region can also respond to non-value related processes including information coding or salience effects (O’Doherty, 2014). To complement our initial neural results, in the preregistered Study 2 we examined a behavioral measure of the value associated with rating expectancy-confirming targets. Specifically, we tested how much money participants were willing to forgo to view stereotype-consistent instead of stereotype-violating targets.

Participants on Amazon Mechanical Turk (*n*s = 174 and 169 in Study 2a and 2b, respectively) made a series of choices to rate one of two target types: a stereotype-consistent target (e.g., a man who enjoys riding motorcycles) or a stereotype-violating target (e.g., a man who enjoys shopping for shoes), designated as “typical” and “atypical” respectively. After each choice, participants saw a target accompanied by a statement and rated the likelihood that the target would be associated with the statement on a 0–100 scale. Participants had up to 5 seconds for each phase of the task (see Fig S2). To avoid potentially different responses to male and female targets, Study 2a included only male faces and Study 2b included only female faces. On each of the 25 trials, small monetary payoffs ($0.03–$0.09 in increments of 2 cents) were associated with each target choice. Participants received a subset of these payoffs as a monetary bonus for the task. Payoff amounts for each target choice varied across trials (and were occasionally equal), as did the location of the option for which participants received the larger amount. If consistency with stereotypes is intrinsically rewarding, participants should be willing to forgo money—to choose the lower-paying option—to see stereotype-consistent individuals. On the other hand, a participant seeking to maximize monetary payoff should consistently choose the higher paying option regardless of the stereotypicality of the information that follows.

We modeled the relative value of each target type by calculating the point of subjective equivalence (PSE) between the two options. This value was derived by fitting a cumulative normal distribution curve to participants’ choices (Fig. 3A) and finding the monetary value at which participants effectively chose arbitrarily between the two target types (Moscatelli *et al.*, 2012). Thus, the PSE represents the relative monetary value of one target type over another.

**Figure 3.**
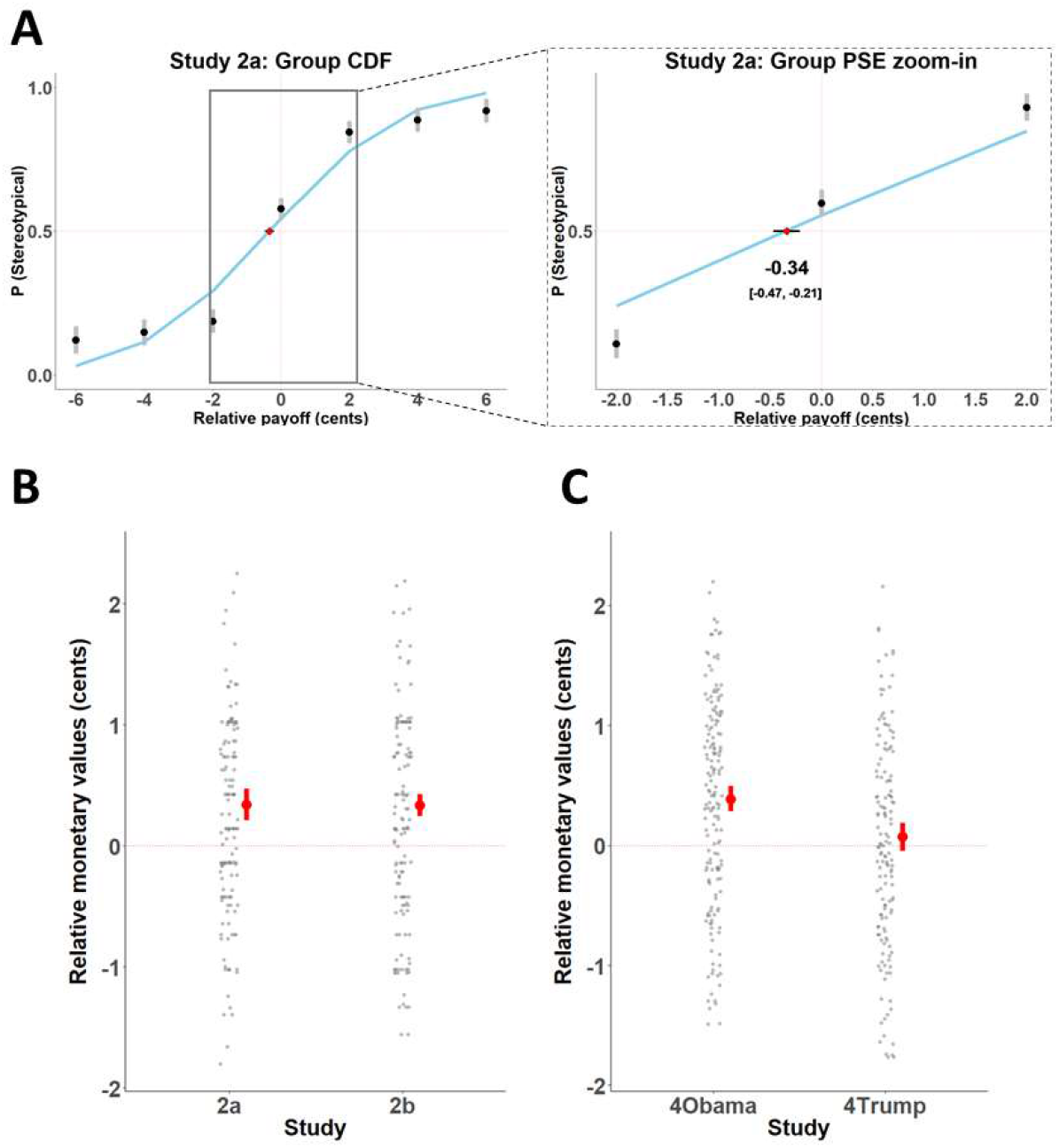
The monetary values of consistency with stereotype and person-specific expectations. (A) Visualization of the cumulative distribution function we used to calculate the Point of Subjective Equivalence (PSE), illustrated by group data from Study 2a. The x-axis represents the difference between the monetary values associated with the two target types presented in each trial. Each dot indicates the proportion of trials in which participants chose to rate a stereotype-confirming over a stereotype-violating target. The PSE was calculated as the point at which a cumulative normal distribution function, fit to these responses, passes 50%. This point represents the relative monetary value associated with one target type over the other. Negative values indicate that participants preferred to incur a relative monetary loss to rate a confirming target. Error bars depict 95% confidence intervals. (B) Distribution of individual PSE values for Study 2. Rating stereotype-consistent targets in Studies 2a and 2b was associated with significantly higher subjective value than rating stereotype-violating targets. Each gray dot depicts PSE for a specific participant. Red dots indicate the sample mean. Error bars depict 95% confidence intervals. (C) Distribution of individual PSE values for Study 4. Rating Obama-consistent trials was associated with significantly higher subjective value than rating Obama-violating trials. However, rating Trump-consistent trials did not significantly differ from rating Trump-violating trials.

As predicted, participants demonstrated a significant preference for seeing stereotype-consistent targets over stereotype-violating targets. When the two trial types shared the same payoff amounts, participants chose the stereotypical targets 58% and 57% of the time for male and female targets, respectively (significantly more than chance, as indicated in a generalized mixed model analysis by an odds ratio of 1.4, *Z* = 3.11, *p* = 0.0019 and an odds ratio of 1.41, *Z* = 2.91, *p* = 0.00369 for the zero-centered intercept in Study 2a and 2b, respectively).

Moreover, the calculated PSE indicated that participants forewent an average of 0.34 and 0.335 cents per trial to rate a stereotype-consistent over a stereotype-violating target in Study 2a and 2b, respectively (95% Confidence Interval [CI]: 0.21–0.47, *t*_(173)_ = 5.17, *p* < 0.0001, *Cohen’s d* = 0.39 [0.24–0.55] and CI: 0.25–0.43, *t*_(168)_ = 7.3, *p* < 0.0001, *d* = 0.56 [0.4–0.72] for the two studies, respectively; Fig 3b). This PSE resulted in an average loss of 10% of potential earnings, as participants chose lower monetary amounts to view stereotype-consistent targets. Just as non-human primates prefer to view dominant groupmates over receiving juice (Deaner *et al.*, 2005) and students are willing to forgo money to talk about themselves (Tamir and Mitchell, 2012) or to view attractive members of the opposite sex (Hayden *et al.*, 2007), our participants gave up money to view information that was in line with their stereotypical expectations.

### Study 3: Neural response to interpersonal expectations

Together, Studies 1 and 2 suggest that perceivers experience consistency with social expectations as intrinsically rewarding. However, we designed these studies primarily to test the effects of a specific type of social expectations – gender-based stereotypes. Although stereotypes are a significant source of interpersonal expectations, perceivers routinely make use of additional, idiosyncratic sources of information, especially for individuals with whom they are highly familiar. Does the reward value of expectancy-consistent information extend to person-specific predictions?

To examine this question, Study 3 assessed the responses of the neural reward system to the consistency with and violation of expectations regarding two highly familiar targets, the current and previous presidents of the United States at the time of the study: Donald Trump and Barack Obama, respectively. To facilitate comparison to Study 1, the preregistered Study 3 also included stereotype-derived expectations about unfamiliar targets. As in Study 1, participants (*n* = 30) rated how likely each of 240 specific statements described a specific man or woman or described Donald Trump or Barack Obama. We presented stereotype-related and person-specific statements in blocks of 15 trials per content domain for a total of 60 trials per condition (consistent/violating targets in the stereotype/person-specific domain). Person-specific statements included various characteristics typically associated with one – but not the other – leader (e.g., “Supports a wall along the borders” versus “Acts to support women’s rights”). Participants first read a short statement for 1.5 seconds and then saw the face of a target (a man, a woman, Trump or Obama; see Fig S1C). Participants rated how likely the statement was to describe the target (see Table S3 and SI Results for full details).

We conducted two parallel analyses to examine whether the NAcc was more engaged when presented with information that was consistent expectations than with information that violated them. First, a whole-brain random-effects contrast identified regions that were more active for *expectation-consistent > expectation-violating* trials (p < 0.05, corrected; see Table S3 for full results). This analysis indicated significantly greater response in several regions, including the NAcc, when the target was consistent with the expectation set by the preceding statement than when the target violated that expectation (Fig. 1B).

In a second analysis we defined the NAcc via two independent procedures, first as bilateral spheres around peak voxels identified in a meta-analysis, and then by examining participants’ neural responses during the MID task. Across both procedures the NAcc demonstrated a robust difference in activity between expectation-consistent and expectation-violating targets (Fig. 1D). However, we did not observe any difference between stereotypes and person-specific trials in any of the analyses. Specifically, a 2 × 2 repeated measures ANOVA over activity in bilateral NAcc revealed a main effect of expectation-consistency, with higher activation associated with expectation-consistent compared to expectation-violation (meta-analysis ROIs: *F*_(1,29)_ = 21.89, *p* < 0.0001, *η^2^_p_* = 0.43 [0.19–0.58]; MID ROIs: *F*_(1,29)_ = 11.2, *p* = 0.002, *η^2^_p_* = 0.28 [0.07–0.46]). Activation did not significantly differ between stereotype and person-specific content (meta-analysis ROIs: *F*_(1,29)_ = 3.64, *p* = 0.07, *η^2^_p_* = 0.11 [0–0.29]; MID ROIs: *F*_(1,29)_ = 1.41, *p* = 0.24, *η^2^_p_* = 0.05 [0–0.2]) and no interaction was observed between the factors (meta-analysis ROIs: *F*_(1,29)_ = 0.31, *p* = 0.58, *η^2^_p_* = 0.01 [0–0.13]; MID ROIs: *F*_(1,29)_ < 0.01, *p* = 0.99, *η^2^_p_* < 0.0001). Accordingly, expectation-consistency yielded more neural activity for each of the two content domains when examined separately (meta-analysis ROIs: *F*_(1,54.8)_ = 5.96, *p* = 0.0179, *η^2^_p_* = 0.1 [0.01–0.23] and *F*_(1,54.8)_ = 11.03, *p* = 0.0016, *η^2^_p_* = 0.17 [0.04–0.31] for gender stereotypes and person-specific expectations, respectively; MID ROIs: *F*_(1,55.47)_ = 4.36, *p* = 0.04, *η^2^_p_* = 0.07 [0.001–0.2] and *F*_(1,55.47)_ = 4.45, *p* = 0.04, *η^2^_p_* = 0.07 [0.002–0.2], respectively). Thus, consistency with social expectations triggered more activation in the NAcc than violation of such expectations, regardless of whether source of the expectation was general social knowledge (stereotypes) or person-specific knowledge.

Finally, similar to Study 1, variability in behavioral responses in Study 3 enabled us to test the effects of subjective endorsement of characteristics for stereotype-based characteristics (but not for person-specific-expectations; see SI results). As in Study 1, NAcc activity was not modulated by subjective endorsement of characteristics for stereotype-consistent targets (simple effects analyses: meta-analysis ROIs: *F*_(1,58)_ = 3.99, *p* = 0.0503; *η_p_^2^* = 0.06 [0, 0.18]; MID ROIs: *F*_(1,56.5)_ = 3.20, *p* = 0.08; *η_p_^2^* = 0.05 [0, 0.17]). However, we did not observe an interaction between response and condition (meta-analysis ROIs: *F*_(1,29)_ = 0.08, *p* > 0.5, *η_p_^2^* = 0.003 [0, 0.09]; MID ROIs: *F*_(1,29)_ = 0.40, *p* > 0.5, *η_p_^2^* = 0.01 [0, 0.14]). In other words, unlike in Study 1, NAcc activation was *in*sensitive to the degree to which a participant judges information to apply to a target, regardless of the stereotypicality of the target. Together, these results suggest that consistency with expectations about other targets is valuable for the two most dominant sources of interpersonal expectations – group-based stereotypes and person-specific knowledge.

### Study 4: The monetary value of person-specific expectation-confirmation

In Study 2, we observed that perceivers are willing to forgo money to view stereotype-consistent (rather than stereotype-violating) information. To examine whether this behavioral effect extends to expectations about specific individuals, Study 4 replicated the procedure from Study 2 using familiar individuals (Obama and Trump). On each of 32 trials, participants on Prolific Academic (n = 189 and 172 in Study 4a and 4b, respectively) first chose between seeing either an expectation-consistent or an expectation-violating target and then rated the target. Study 4a included only Obama as the target of statements and Study 4b included only Trump. After choosing the type of content they would like to see, participants rated the likelihood that a specific statement would be associated with the target on a 0–100 scale. Payoff amounts for each choice varied across trials (and were occasionally equal), as did the option for which participants received the larger amount. As in Study 2, we quantified the subjective monetary value of each option by calculating the PSE between the two display types by fitting a cumulative normal distribution curve to participants’ choices and finding the monetary value at which participants were indifferent to the two options. If participants experience consistency with expectation as equally rewarding regardless of the target of expectations, then they should choose to forgo money to rate statements consistent with Obama and Trump in Studies 4a and 4b, respectively.

The results from Study 4a indicate that participants preferred to see expectation-consistent statements of Barack Obama. When the payoff amounts were equal for confirming and violating statements, participants chose the consistent statements 60% of the time (significantly more than chance, as indicated in a generalized mixed model analysis by an odds ratio of 1.42, *Z* = 4.74, *p* < 0.0001 for the zero-centered intercept in Study 4a). Moreover, the calculated PSE indicated that, on average, participants gave up 0.39 cents per trial to rate an expectancy-consistent over an expectancy-violating statement about Obama (95% [CI]: 0.28–0.49, *t*_(188)_ = 7.21, *p* < 0.0001, *d* = 0.52 [0.37–0.68]; Fig 3C). However, the same was not true for Trump-related statements in Study 4b. At equal payoff amounts, participants had no preference between expectancy-consistent and expectation-violating statements (choosing the consistent option 49% of the time; odds ratio of 1.01, *z* = 0.13, *p* = 0.9), and the calculated PSE was not different from zero (0.07 cents; [−0.05–0.19], *t*_(173)_ = 1.17, *p* = 0.12, *d* = 0.09 [−0.06–0.24]; Fig 3C). The difference between the studies was significant (*t*_(328.29)_ = 2.28, *p* = 0.023, *d* = 0.25 [0.03–0.46]). However, this finding should be interpreted with caution, as our sample was demographically skewed towards liberals, potentially diluting the effect of consistency with expectations from Trump with additional factors (see Fig S4). Together, these findings suggest that the rewarding effect of expectation-consistency is not limited to stereotypes, but also applies to knowledge about specific individuals. Notably, however, not all sources of knowledge equally contribute to the reward value; people were not willing to forgo monetary amounts to rate content consistent with their expectations about Donald Trump (see SI results for potentially related findings in Study 3).

## Discussion

The human preference for consistent and predictable social interactions has long been acknowledged as a core motivational component driving everyday behavior (Festinger, 1957; Gawronski, 2012). To predict the behavior of others, perceivers regularly employ biased strategies to collect and interpret information that corresponds to their expectations (Lord *et al.*, 1979; Johnston and Macrae, 1994; Frimer *et al.*, 2017; Falben *et al.*, 2019; Oyserman and Yan, 2019). Here we provide evidence to suggest that humans associate expectation-consistent information with intrinsic value, much like other forms of reward such as food or money. Our findings suggest that this reward value is generated regardless of the source of the social expectation. Participants were willing to forgo money to rate an expectation-consistent target rather than its expectancy-violating counterpart. Moreover, doing so was associated with increased activity in a brain region in the neural reward circuitry, regardless of whether the target was consistent with gender stereotypes or knowledge about US presidents. Put simply: people find it rewarding to have their expectations (stereotypical or idiosyncratic) confirmed.

This line of research coincides with emerging theories that highlight the instrumental value of behaviors and perceptions that fall in line with our expectations. In an early example of this value, Allport (Allport, 1954) described a mental process in which ‘A Scotsman who is penurious delights us because he vindicates our prejudgment’ (p.22). Some recent theories suggest that, because most of our social expectations are anchored in our social environment, repeated interaction with expectancy-confirming information leads to continuous reinforcement of our expectations (Huebner, 2016; Oyserman and Yan, 2019; Peters, 2020). Once established, these expectations induce motivations and cognitive representations that persist even in the face of disconfirmation (Berridge, 2012; Uusberg *et al.*, 2019; Yon *et al.*, 2019). One theory further suggests that the metabolic costs associated with the violation of expectations increase the desirability of expectation-consistent behavior from an evolutionary standpoint (Theriault *et al.*, 2020). In line with these theories, the current studies demonstrate that information consistent with social expectations is indeed associated with subjective value.

The current findings provide a neural extension to prominent accounts of implicit (i.e., involuntary) stereotyping and prejudice (Tibboel *et al.*, 2015; Greenwald and Lai, 2020). Group-based stereotypes typically draw on categorical distinctions to facilitate easier decision making by enabling faster and more efficient processing of stereotype-confirming information (Roese and Sherman, 2007). Stereotypes are also more familiar, thus allowing perceivers to process such information fluently (e.g., Smith *et al.*, 2006). Downstream, perceivers evaluate expectation-confirming individuals more positively, allocate more economic resources to them, and judge them as more hirable (Phelan and Rudman, 2010; Stern *et al.*, 2015; Stern and Rule, 2018). Complementarily, violations of social expectations pose a threat to individuals and social structures alike, which, in turn, often try to eliminate the threat and reinforce the original expectation (Morgenroth and Ryan, 2020). Our results provide a candidate mechanism for these effects, whereby the preference of expectation-consistent information translates into a subjective value that shapes how we evaluate specific individuals (cf., Amodio and Devine, 2006). Although our findings are mute with respect to the specific mechanism driving the rewarding effect (e.g., a motivational goal to confirm expectations or processing fluency), our results hint at when perceivers can assign value to expectation-violating information. In our task, behavior asymmetrically affected the neural response in the nucleus accumbens. Whereas stereotype-consistent targets always evoked the same level of neural activity regardless of participants’ responses, in Study 1 stereotype-violating targets elicited enhanced NAcc activity only if perceivers judged them as likely to be associated with the expectancy. This pattern suggests that participants can assign value to stereotype-violating targets, perhaps depending on the believability of the counter-stereotypical judgment.

The asymmetric effect of behavior on the neural response in the NAcc also suggests that our findings do not result only from participants’ desire to be correct in their predictions. Previous studies used objectively measurable performance to demonstrated increased ventral striatum activity when participants provided correct responses, either with external feedback (Ullsperger and von Cramon, 2003; Tricomi and Fiez, 2008) or in its absence (Satterthwaite *et al.*, 2012; Ruissen *et al.*, 2018). If the correspondence between the targets and predictions would have been the sole process driving the effects, then behavioral ratings of targets would not have interacted with types of prediction to modulate NAcc activity. Nonetheless, future studies should experimentally control for this alternative interpretation by, for example, including an explicit prediction phase before presenting a target and yoking the identity of the target to participant’s prediction.

The expectation-consistency account of ventral striatum activity we put forward complements earlier hypotheses that activity in the mesolimbic circuit reflects prediction errors (Daw *et al.*, 2011). Prediction errors refer to discrepancies between actual and expected outcomes, typically in the context of learning (Niv and Schoenbaum, 2008). Numerous studies have found that the ventral striatum increases its activity as the gap between the expected and the received outcome grows (e.g., Glimcher, 2011; Ballard *et al.*, 2018). However, the present results diverge from this prediction error pattern. We observed increased striatal activity when an outcome—the target face—was not different from the expected outcome; a prediction error account would suggest that such activity should accompany unexpected information. Notably, unlike the vast majority of studies exploring the prediction error account, the paradigms we used in the current investigation did not involve learning. Participants saw each target only once during the entire study and formed their expectations based solely on previously established knowledge.

Furthermore, our paradigm did not involve any explicit feedback. Therefore, participants had no objective external verification of their predictions, nor did we directly measure their predictions. Future studies could formally test whether striatal activity in response to expectancy-consistent information also emerges when participants experience feedback and need to learn the new information.

The current set of studies tested the value of expectancy-consistency through relatively innocuous stereotypes and associations rather than overtly positive or negative bits of information (with one potential exception of Trump-related statements for liberal participants; see below). Including only neutral or mildly-valenced statements allowed us to test the direct effects of expectation-confirmation with little influence from potentially competing motivations, such as social desirability. Therefore, we cannot determine whether perceivers will continue to value stereotype-confirming information even when they are motivated to suppress them. Our paradigms provided only indirect evidence on this question. In Study 3, our sample consisted of liberal participants (average rating of 3.37 on a 1-9 scale, ‘1’ denoting extremely liberal and ‘9’ denoting extremely conservative) for whom information consistent with Donald Trump might be aversive. However, these participants demonstrated comparable activation in the neural reward circuitry in response to statements about Donald Trump and Barack Obama, implying that seeing expectation-confirming information is rewarding regardless of the valence of these statements. In Study 4, conversely, participants (mean difference of 64 points on a 0-100 scale in favor of liking Barack Obama; see SI Results) did not choose to incur a cost to see Trump-consisting information, suggesting that additional motivations affected their behavior. These first steps warrant further research to characterize the precise mechanisms contributing to subjective value perceivers attribute to confirmation of expectation and the subsequent behaviors associated with this value.

An additional and important future route of investigation should explore the generalizability of ou r findings to non-social contexts. Here we focused on expectations concerning social targets—unfamiliar men and women as well as familiar targets. We cannot ascertain whether similar effects would emerge for non-social expectations, such as about inanimate objects or the weather. Given some accounts suggesting that social information processing has a preferred status in the primate brain (Adolphs, 2009; Atzil *et al.*, 2018; Lockwood *et al.*, 2020), we might expect that (a) the subjective value attributed to consistency should be greater for the kinds of social expectations studied here and (b) confirmation of *social* expectations would lead to responses similar to reward-like responses for primary reinforcers like food.

Altogether, these findings join a growing body of literature that characterizes how our prior beliefs modulate information-processing to fortify a world view and protect established expectations (Golman *et al.*, 2017; Charpentier *et al.*, 2018; Jepma *et al.*, 2018; Gershman, 2019; Yon *et al.*, 2019). We suggest that the subjective value imbued upon targets who conform to societal expectations may serve to sustain multiple stereotype- and expectation-induced biases. To mitigate the negative implications associated with these expectations, society will need to acknowledge the subjective value associated with their confirmation.

## Materials and Method

### Study Design

#### Materials

To create expectation-setting statements we generated a list of verbal statements that described relevant individual preferences, traits, behaviors or professions. Studies 1 and 2 included 136 gender-related and 68 gender-neutral statements (see SI Materials and Methods). We verified the stereotypicality of these statements in a pilot study (*n* = 78) in which participants from the local community indicated how typical the characteristic was for a specific gender on a visual scale of 0 (”very untypical”) to 100 (”very typical”; the scale had no other tick marks). Each participant was randomly assigned to rate each statement either for men or for women. Participants were instructed to base their ratings on how they thought the average person would respond. This verification procedure was successful; men were associated with men-stereotypic statements more than women (mean difference: 25.7) and women were associated with women-stereotypic statements more than men (mean difference: 25.1). Overall, the statements contained 2-9 words (mean: 4.69, S.D: 1.44; no difference between experimental conditions, *p* > 0.2) and 9-45 characters (mean: 26.68, S.D.: 7.69; *p* > 0.18). Studies 3 and 4 further included 120 person-specific statements pertaining to Barack Obama and Donald Trump. A total of 243 participants from Amazon Mechanical Turk rated a sample of 60 of these statements, randomly determined per participant. On each trial participants indicated how typical the presented characteristic was for the two targets (a separate scale for each target; the two scales were presented simultaneously with a randomly determined order). Obama was associated with Obama-related statements more than Trump (mean difference: 56.1) and Trump was associated with Trump-related statements more than Obama (mean difference: 54.5). Overall, the statements contained 2-9 words (mean: 4.65, S.D: 1.52; no difference between experimental conditions, *p* > 0.5) and 11-45 characters (mean: 28.33, S.D.: 8.43; *p* > 0.5).

#### Studies 1 and 3

Twenty-eight individuals participated in Study 1 and 30 individuals participated in Study 3. Additional participants were excluded due to excessive motion, technical issues or lack of response to more than 20% of trials (3 and 6 participants from Studies 1 and 3, respectively). All participants provided informed consent in a manner approved by the Committee on the Use of Human Subjects in Research at Harvard University. Study 3 was pre-registered (https://osf.io/h9c6x/?view_only=6fa03fc0ceb04e2083db9e485dbe6615). See SI for demographic information. The current sample size allowed a power of 0.8 to detect a medium effect size (*Cohen’s d* = 0.5) in the planned one-tailed contrast between expectation-consistent and expectation-violating targets at the region of interest. In both studies participants formed impressions about target individuals. On each trial, participants first saw a statement for 1.5 seconds. The statements in Study 1 described a stereotypically neutral, stereotypically male, or stereotypically female characteristic; statements in Study 3 described a characteristic which was either stereotypically male or female or closely associated with Barack Obama or Donald Trump (see https://osf.io/tgja3/?view_only=d849bad1b606474f80c1a3e40e740875 for open materials). Next, participants saw the statement with a face of a man or a woman (in Study 1; Study 3 also included face images of the relevant leaders). The statement-face pair appeared on screen for 4 additional seconds (3.5 seconds in Study 3) for a total of 5.5 seconds (5 seconds) per trial. Each trial ended with a 0.5 second fixation crosshair. Participants used their left hand to indicate how likely the presented target was to be described by the specific characteristic using a 4-point scale (1-‘very unlikely’; 4 -‘very likely’). Participants responded while the pair appeared on screen. Trials in both studies were separated by variable intertrial intervals of 0-9s (Dale, 1999) optimized for our contrast of interest (see SI Materials and Methods for details).

#### Studies 2 and 4

A total of 343 participants were included in Study 2 (174 in Study 2a) and 361 in Study 4 (189 in Study 4a). Additional participants were excluded by criteria set in the pre-registered protocols for each study (see SI Materials and Methods for details; see also https://aspredicted.org/blind.php?x=rt3wi6 (Study 2a), https://aspredicted.org/blind.php?x=db3gz4 (Study 2b), and https://aspredicted.org/blind.php?x=9z83yb (Studies 4a and 4b)). Sample size was set to allow sufficient power (0.8) to detect a small effect size (*Cohen’s d* = 0.2) in a one-sample t-test for each study. Informed consent was obtained from all participants in a manner approved by the Committee on the Use of Human Subjects at Harvard University.

### Statistical Analysis

#### Studies 1 and 3

To localize brain regions associated with the processing of rewarding stimuli, we defined 8-mm spheres around peak coordinates drawn from a comprehensive meta-analysis (Bartra *et al.*, 2013). To functionally identify these brain regions, participants in both studies completed a Monetary Incentive Delay (MID) task (Knutson *et al.*, 2000) immediately after the impression formation task (see Fig. S1B and SI Materials and Methods for details). This task allows the identification of monetary-reward-sensitive ROIs by comparing trials in which participants won money to trials in which participants could not earn any reward. We extracted and averaged parameter estimates across voxels in each ROI per condition of interest and analyzed them using within-participant ANOVAs as implemented by afex package (Singmann *et al.*, 2018) for R, version 0.22-1. We plotted the results using the package ggstatsplot (Patil, 2018), version 0.2.0. Additionally, we plotted the mean effect size and the bootstrapped 95% confidence intervals using the package dabestr (Ho *et al.*, 2019), version 0.2.2.

We collected neuroimaging data with a 3T Siemens Prisma scanner system (Siemens Medical Systems, Erlangen, Germany). First, we acquired high-resolution anatomical images using a T1-weighted 3D MPRAGE sequence. Next, whole brain functional images were collected using a simultaneous multi-slice (multiband) T2*-weighted gradient echo sequence (TR = 2000 msec, TE = 30 msec, voxel size = 2 × 2 × 2 mm3, 75 slices auto-aligned to −25 degrees of the AC-PC line). Participants completed four impression formation task runs consisting of 229 volumes each (245 volumes in Study 3). Finally, participants completed the MID task in a single run consisting of 110 volumes using identical parameters to those mentioned above. We used SPM12 version 6225 (Wellcome Department of Cognitive Neurology, London, UK) to process and analyze the fMRI data. Data were corrected for differences in acquisition time between slices, corrected for inhomogeneities in the magnetic field using fieldmap (Cusack and Papadakis, 2002), realigned to the first image to correct for head movement, unwarped to account for residual movement-related variance and co-registered with each participant’s anatomical data. Functional data were then transformed into a standard anatomical space (2 mm isotropic voxels) based on the ICBM152 brain template (Montreal Neurological Institute). Normalized data were then spatially smoothed (6 mm full-width at half-maximum, FWHM) using a Gaussian Kernel (see SI Materials and Methods for full details of scanning and analysis procedures). We analyzed preprocessed data using a general linear model in which we modeled trials as boxcar functions with an onset at face presentation (1.5s after statement presentation) and with variable duration determined per trial by reaction time to control for effects of reaction time on the neural response (Grinband *et al.*, 2008). Our main analysis included a model in which we conditionalized trials based on trial type (stereotype-consistent, stereotype-neutral or stereotype-violating trials in Study 1; stereotype-consistent, stereotype-violating, person-specific-confirming and person-specific-violating trials in Study 3). In our secondary analysis (Fig 2) we split each of the trial types included in the main analysis with two regressors, one in which participants provided a “Likely” rating and one in which they provided an “Unlikely” rating. We convolved events with a canonical hemodynamic response function and its temporal derivative and included additional covariates of no interest (session mean, no response trials, six motion parameters and their temporal derivative). The final first-level GLM was high-pass filtered at 128 s. Analyses were performed individually for each participant, and contrast images were subsequently entered into a second-level analysis treating participants as a random effect. We report activations that survived a threshold of p<0.001 (uncorrected) at the voxel level and (cluster-size) corrected to p<0.05 at the cluster level using Monte Carlo simulations (1,000 iterations) with the current imaging and analysis parameters (Slotnick, 2017).

#### Studies 2 and 4

We analyzed choice data with logit generalized linear mixed models as implemented in the lme4 package version 1.1-14 (Bates *et al.*, 2014) for R version 3.4.2 (R Core Team, 2017). We used the probit link and included fixed effects for the intercept and for the value difference between the two target types, as well as random effects for the intercepts for participants for the by-participant random slopes for the fixed effect of value difference. We calculated the PSE (and the related SEs) for the difference between the two target types for each study by the Delta Method as implemented in the MixedPsy package (Moscatelli *et al.*, 2012; Moscatelli and Balestrucci, 2017) for R. The Delta Method relies on responses to <u>all trials</u> aggregated across participants in a generalized linear model to approximate the PSE with a Gaussian distribution (Moscatelli *et al.*, 2012) and to plot the cumulative distribution function. The model included the difference between the two target types on each trial as the predictor value and a binary outcome (stereotypical/knowledge-consistent option chosen) as the predicted value, with the probit link. To present individual-level data (Figure 3B and 3C), we calculated the PSE for each participant. This resulted in some PSE values that exceeded the possible values in the current studies, as PSE models do not accurately reflect behavior when one choice option is rarely selected, as was the case for some participants. Therefore, we excluded participants for whom the calculated PSE value exceeded ±6 (*n* = 4–14 across studies).

## Acknowledgements

This work was partially supported by the Israeli Science Foundation (No. 79/18, to N.R.). N.R was supported by Yad Ha’Nadiv (Rothschild) Foundation and the Mind, Brain and Behavior interfaculty program at Harvard University. Parts of this work were conducted at the Harvard Center for Brain Science, which is supported by the NIH Shared Instrumentation Grant Program (No. S10OD020039). We thank Kirstan Brodie for her assistance with data collection for Study 3.

## Supplementary Materials

**Supplementary Materials include:**

Supplementary text

Figures S1 to S4

Tables S1 to S6

SI References

### Supplementary Materials: Materials and Methods

Below we provide a detailed description of all the materials and methods used including all measures and data exclusions.

#### Participants

##### Studies 1 and 3

Twenty-eight individuals participated in Study 1 (mean age: 21.85, S.D.: 2.95, range: 18–30, 21 females, 15 Caucasian, 4 Hispanic, 4 mixed, 3 Asian, 2 African American) and 30 individuals participated in Study 3 (mean age: 22.37, S.D.: 2.62, range: 18–30, 17 females, 14 Caucasian, 7 Asian, 4 African American, 3 mixed, 1 Hispanic, 1 did not self-identify). Participants were recruited from Harvard University and its surroundings using Harvard’s Psychology Study Pool website. Additional participants were excluded due to technical issues (1 participant from Study 3), lack of response to more than 20% of trials (2 participants from each study) or excessive movement (more than 1mm – half-width of the acquired voxel size) in more than one functional run in the scanner (1 participant in Study 1, 3 participants in Study 3). All participants were healthy, right-handed, native English speakers with normal or corrected-to-normal vision and no history of neurological or psychiatric conditions. Participants were compensated with $65 in Study 1 and $55 in Study 3. We determined supplemental compensation (up to $30) based on participants’ performance in the monetary incentive delay task (see below). Pilot versions of the imaging tasks were conducted outside the scanner to ensure the functionality of the task.

##### Studies 2 and 4

Three-hundred-ninety-six individuals from Amazon Mechanical Turk completed Study 2 (in Study 2a, 198 out of 216 individuals who started the study; in Study 2b, 198 out of 223 individuals). Study 2a was conducted in March 2018, and Study 2b was conducted in April 2018. Four-hundred-seventy individuals from Prolific Academic completed Studies 4a and 4b in January 2019 (235 in each study; a total of 503 participants started the study). All participants were 18 years old or older, had an approval rate of 95% or higher, held a US nationality and participated only in one of the studies. In addition, participants from Amazon Mechanical Turk were required to have completed at least 100 tasks to be eligible for Study 2. The final sample size we report in the main text incorporates the following pre-registered exclusion criteria for participants who (1) answered incorrectly at least two of the manipulation or attention checks, (2) completed the survey in more than 20 minutes or less than 2.5 standard deviations below the sample mean duration, (3) failed to answer more than 15% of the survey, (4) provided similar ratings (on the 0–100 scale) on all trials; specifically, participants whose standard variation of the rating was 2 standard deviations below the sample mean standard deviation (not applied in Study 2a), (5) provided an identical response in all 2AFC trials (applied only in Study 4) and (6) indicated in their debriefing that they had understood the goal of the task and had acted per this understanding at any point during the task. Participants in all studies were similarly distributed on gender (Study 2a: 51.72% males, 45.98% females; Study 2b: 50.3% males, 48.52% females; Study 4a: 51.32% males, 46.56% females; Study 4b: 41.86% males, 56.98% females; across studies 1%–3% of participants self-identified as gender-nonconforming or other identification), age (Study 2a: Mean [S.D.]: 37.47 [11.75]; Study 2b: 35.13 [10.04]; Study 4a: 31.22 [10.32]; Study 4b: 31.99 [10.97]) and ethnicity (Study 2a: 78.16% White/European American; Study 2b: 74.56% White/European American; Study 4a: 69.84% White/European American; Study 4b: 69.19% White/European American). Study 4 measured political orientation on a 1–9 scale (see below). We observed no differences on this measure between Study 4a and Study 4b (Study 4a: 3.79 [2.13], Study 4b: 3.45 [1.96]).

#### Stimuli

All studies included face-statement pairs. We selected faces from the 10k US Adult Faces Database (a large-scale database of natural face photographs of the U.S. adult population (Bainbridge *et al.*, 2013)). The current investigation focused on Caucasian faces to avoid intrusion of racial or intersectional stereotypes. We further restricted the faces to be moderately memorable (0.4 – 0.6 hit rate in the 10k database, computed over an average of 81.7 people per photo) and excluded faces with any kind of distinguishable accessories (e.g. hats, big necklaces, etc.). This resulted in a total of 204 faces, 102 per gender. Three additional photos per gender were used in the practice sessions. All face pictures had neutral to mildly positive expressions. We resized all pictures to 240 by 256 pixels and presented them in color on a gray background (see Fig S1). For Studies 3 and 4 we also included images portraying the faces of Barack Obama and Donald Trump (a single image per leader). These images were taken from open access sources and cropped to be identical in dimensions to the rest of the face stimuli.

We generated gender-stereotype-relevant statements in the form of individual preference, trait, behavior or profession (e.g., “Loves taking risks”, “Doesn’t cry”, “Swears a lot”, “Is a truck driver”, etc.; see https://osf.io/tgja3/?view_only=d849bad1b606474f80c1a3e40e740875). We piloted the verbal statements in a two-phase procedure using non-overlapping groups of participants that did not participate in the reported studies. In the first session, 23 participants rated 160 gender-related statements generated by the authors based on various sources (e.g., Prentice and Carranza, 2002). Participants saw each statement once and indicated how typical the characteristic was for a specific gender on a visual scale of 0 (= “very untypical”) to 100 (= “very typical”; the scale had no other tick marks). Each participant rated each statement either for men or for women, never for both, randomly determined per participant. Participants were instructed to base their ratings on how they thought the average person would respond. Based on the ratings from this first phase, we selected 136 gender-related statements (68 per gender) for which the average rating was at least 65 for one of the genders and less than 35 for the other gender. In a second session of the pilot testing, we verified the stereotypicality of these statements with a total of 78 participants who used the same rating procedure to rate the selected 136 statements and an additional 68 gender-neutral statements (e.g., “Exercises regularly”, “Thinks positively”, “Is a TV reporter”). The statements included in the studies contained 2–9 words (mean: 4.69, S.D: 1.44; no difference between experimental conditions, *p* > 0.2) and 9–45 characters (mean: 26.68, S.D.: 7.69; *p* > 0.18).

We utilized a similar procedure for leader-relevant statements. First, we generated 160 statements based on common knowledge about the selected leaders. We constructed the statements such that a statement consistent with knowledge about one leader would violate knowledge about the other leader. We piloted these statements with 243 Amazon Mechanical Turk participants. Each participant saw a randomly selected subset of 40 or 60 statements and rated them on two scales: “how likely is the statement to be attributed to Barack Obama” and “how likely is the statement to be attributed to Donald Trump” on a visual scale of 0 (= “very unlikely”) to 100 (= “very likely”). Participants were instructed to base their ratings on how they thought the average person would respond. Each statement had an average of 81.75 raters (S.D.: 6.23; range: 67–100). We selected the 120 statements that differed the most between ratings for the two leaders. Overall, the statements contained 2–9 words (mean: 4.65, S.D: 1.52; no difference between experimental conditions, *p* > 0.5) and 11–45 characters (mean: 28.33, S.D.: 8.43; *p* > 0.5).

#### Behavioral procedure

##### Studies 1 and 3

The impression formation task included 204 face-statement pairs in Study 1 and 240 face-statement pairs in Study 3. To create stereotype-consistent and stereotype-violating pairs, we yoked half of the 136 stereotypical statements (120 statements in Study 3) with faces from the gender corresponding to the stereotype and the other half with faces from the mismatching gender (yoking of statements was randomized across participants; e.g., the statement “Doesn’t cry” could be paired with a male face for one participant and with a female face for another participant to create a stereotype-consistent or a stereotype–violating pair, respectively). Gender-neutral statements (presented only in Study 1; 68 statements) were randomly and evenly yoked with male and female faces. We randomized the specific face identity paired with each statement across participants. We used a similar procedure to create leader-specific pairs (60 statements per leader), with the exception that a single photo was used per leader.

###### Impression Formation Task

Each trial in the Impression Formation task (see Fig S1) started with a statement (describing a neutral, stereotypically male or stereotypically female characteristic in Study 1; stereotypically male, stereotypically female, Obama-specific or Trump-specific in Study 3) presented for 1.5 seconds. Then, a face joined the statement to form a face-statement pair that was either consistent or inconsistent (or, in Study 1, neutral) with gender stereotypes or (in Study 3) person-specific knowledge. The pair appeared on screen for 4 seconds for a total of 5.5 seconds per trial in Study 1, and 3.5 seconds for a total of 5 seconds in Study 3. Each trial ended with a 0.5-second fixation crosshair. For each pair, participants used their left hand to indicate how likely the presented target was to be described by the specific characteristic using a 4-point scale (1-‘very unlikely’; 4 -‘very likely’). Participants were to provide a response while the pair appeared on screen.

Each of four functional runs included 51 (in Study 1) or 60 (in Study 3) unique face-statement pairs (17 per condition in Study 1, 15 per condition in Study 3). In Study 3 we presented statements in mini-blocks by knowledge domain (stereotypes versus person-specific), 2 blocks per type per run; each mini-block contained 15 trials. Order of the mini-blocks was randomly determined within each run with a limitation that consecutive blocks never displayed the same type of content. To optimize estimation of the event-related fMRI response, we intermixed conditions in a pseudorandom order and separated trials by a variable interstimulus interval (Study1: 0–9 seconds, mean: 3.03, S.D.: 2.25; Study 3: 0–7 seconds, mean: 1.24, S.D: 1.84). We used OptSeq2 (Dale, 1999) to generate sequences optimized for the efficiency of a 3-conditions design for a first-order counterbalanced event sequence in Study 1 and for the efficiency of the (expectancy-consistent versus expectancy-violating) contrast for a first-order counterbalanced event sequence in Study 3. Of these sequences, we selected 6 sequences that contained no more than 5 consecutive events of the same condition (separate sequences were generated and selected per Study). We randomly assigned (with replacement) an event sequence for each functional run to avoid spurious results attributable to differences between conditions in one specific event sequence (Mumford *et al.*, 2014). Within conditions, trials were presented in random order. To facilitate familiarization with the task, participants completed a brief practice session before entering the MRI machine. This practice session included six statement-face pairs in Study 1 and 12 pairs in Study 3; the stimuli used in the practice were not used in any other phase of the experiment.

###### Monetary Incentive Delay (MID) Task

After completing the impression formation task, participants in Studies 1 and 3 completed the Monetary-Incentive Delay (MID) task (Knutson *et al.*, 2000) to allow us to localize brain regions associated with the processing of rewarding stimuli in a non-social context. Participants were not informed about this task before its execution to prevent them from forming an association between the main task and reward processing.

The MID task included a series of trials in which participants attempted to respond, via a button press, to a briefly presented target (a white rectangle) (see Fig S1B). Each trial started with a cue (a blue circle or a green circle) that was presented for 0.5 seconds. The green circle predicted a modest monetary reward ($1) upon a successful response to the target, whereas the blue circle predicted no reward. Nevertheless, participants were instructed to respond to both cue types. Cues were followed by a delay interval randomly varying in duration between 2 and 2.5 seconds. The target was then briefly presented for a duration varied between 130 and 350 milliseconds. Duration varied as a function of the participants’ performance. Specifically, we implemented a 2-down 1-up staircase procedure to create a level of difficulty that would allow participants to successfully respond to the target on two-thirds of the trials. This algorithm succeeded; on average, participants were rewarded on approximately 20 of the 30 trials (Study 1 mean: 20.26; Study 3: 20.21). At the end of each trial, participants saw the amount of money they had earned on that trial along with the total amount they had earned during the task up to that point (presented for 0.5 seconds). The task included 45 trials (with 30 green cue trials). We added participants’ gains in this task to their overall compensation.

###### Memory Test

Once outside the scanner, participants completed a surprise associative memory task that will be reported elsewhere (Reggev & Mitchell, in preparation). Briefly, none of the results reported in the current manuscript were affected by including memory in the analyses.

###### Additional measures

Next, participants completed several individual differences and explicit attitudes scales measuring beliefs about sexism (Ambivalent Sexism Inventory – ASI) (Glick and Fiske, 1996), social dominance orientation (SDO) (Ho *et al.*, 2015), motivation to control sexism (MCS) (Klonis *et al.*, 2005) and need for cognitive closure (NFC) (Kruglanski *et al.*, 1993). Order of the scales was randomized between participants. We included these scales to facilitate future individual differences analyses. Individual differences scores were not used in the current manuscript in any of the analyses due to insufficient power and are reported solely for full disclosure’s sake.

Finally, participants in Study 1 indicated the extent to which they thought the different statements presented during the impression formation task were associated with women and men using the procedure used for piloting the statements. We presented each statement with the gender to which it was yoked in the impression formation task. Participants had up to 10 seconds per trial and were told that they should base their judgments on their own beliefs about men and women, rather than on what the “average” person in the population thinks.

After completing these tasks, participants provided demographic details (age, self-identified gender and self-identified race in an open response format, and in Study 3 political affiliation on a 1–9 scale, 1 = “extremely liberal” and 9 = “extremely conservative”). Then, we probed participants for their understanding of the goal of the study and asked whether they had suspected a memory test. Lastly, we paid and fully debriefed them.

#### Imaging procedure

Images were collected with a 3T Siemens Prisma scanner system (Siemens Medical Systems, Erlangen, Germany) using a 64-channel radiofrequency head coil. Stimuli were projected onto a screen at the end of the magnet bore that participants viewed via a mirror mounted on the head coil. Stimulus presentation was controlled by PsychoPy v1.84.2(Peirce, 2007) running under Windows 7. Prior to entering the scanner participants were extensively briefed by one of the authors about potential movements that can occur in the scanner and ways to mitigate them. Participants were then set up in the scanner, head first and supine in the scanner bore, with a response box in their left hand. Foam cushions were placed within the head coil to minimize head movements. First, high-resolution anatomical images were acquired using a T1-weighted 3D MPRAGE sequence (TR = 2200 msec, TI = 1100 msec, acquisition matrix = 256 × 256 × 176, flip angle = 7, voxel size = 1 × 1 × 1 mm^3^). Second, a fieldmap was acquired in the same plane as the functional images (see below) to correct for inhomogeneities in the magnetic field (Cusack and Papadakis, 2002). Next, whole-brain functional images were collected using a simultaneous multi-slice (multiband) T2*-weighted gradient echo sequence, sensitive to BOLD contrast, developed at the Center for Magnetic Resonance Research (CMRR) at University of Minnesota (Feinberg *et al.*, 2010; Moeller *et al.*, 2010; Xu *et al.*, 2013) (TR = 2000 msec, TE = 30 msec, voxel size = 2 × 2 × 2 mm^3^, 75 slices auto-aligned to −25 degrees of the AC-PC line, image matrix = 104 × 104, FOV = 208 * 208 mm^2^, flip angle = 75°, GRAPPA acceleration factor = 2, multiband factor = 3, phase encoding direction = A -> P). After a brief practice run (identical in content to the practice session completed before entering the scanner), participants completed four impression formation task runs consisting of 229 volumes each in Study 1 and 245 volumes each in Study 3; all runs were complemented by two additional dummy scans and an initial period of approximately 26 s dedicated to references for the GRAPPA procedure. The first four volumes from each run (i.e., in addition to dummy scans) were discarded to ensure T1 equilibrium. The last 5 volumes from each run always included a crosshair fixation to ensure the appropriate estimation of the hemodynamic function for the last events in the run. Finally, participants completed the MID task in a single run consisting of 110 volumes using identical parameters to those mentioned above.

#### Imaging analysis

We processed and analyzed the fMRI data using SPM12 version 6225 (Wellcome Department of Cognitive Neurology, London, UK) on a 2015b MATLAB platform (Mathworks, Natick, MA, USA). Functional data were corrected for differences in acquisition time between slices, corrected for inhomogeneities in the magnetic field using the fieldmap (Cusack and Papadakis, 2002), realigned to the first image to correct for head movement using a 2^nd^ degree B-spline interpolation, unwarped to account for residual movement-related variance using a 4^th^ degree B-spline interpolation and co-registered with each participant’s anatomical data. Then, the functional data were transformed into standard anatomical space (2 mm isotropic voxels) based on the ICBM152 brain template (Montreal Neurological Institute). Normalized data were spatially smoothed (6 mm full-width at half-maximum, FWHM) using a Gaussian Kernel. In addition to the GLM models reported in the main text, in Study 1 we also examined a model in which the stereotypicality of trials was modeled continuously rather than with the dichotomous binning approach. Specifically, this additional model included two regressors – one for trials including a woman’s face and another for trials including a man’s face. We included a separate parametric modulator for each regressor to model the extent of the stereotypicality of the statement included in that trial based on our pilot ratings. For example, the statement “Can lift heavy things” was rated as related more to men than to women in our pilot studies with a 28-points difference on the 0–100 scale. Consequently, the parametric modulation value was 0.28 for trials in which this statement was presented with a man’s face, and −0.28 for trials in which this statement was presented with a woman’s face. Similar to the models reported in the main text, in this additional model we convolved events with a canonical hemodynamic response function and its temporal derivative and included additional covariates of no interest (session mean, no response trials, six motion parameters, and their temporal derivative).

#### Regions of interest (ROIs)

We used two complementary approaches to localize regions involved in the processing of rewards. For independently defined ROIs, we defined 8-mm spheres around peak coordinates drawn from a recent meta-analysis (Bartra *et al.*, 2013). Specifically, we utilized the peaks of the region identified as supporting the processing of both monetary and primary incentives: bilateral ventral striatum (x=−6, y=10, z=−6 and x=10, y=12, z=−6).

To functionally locate these regions, we examined the MID task to identify voxels that responded more to rewarded trials (i.e., trials in which participants successfully responded to the target) than to no-reward trials (i.e., trials in which no reward was available). Whole-brain corrected clusters (using the procedure described in the main text) were defined as independent ROIs.

As we had no prediction about laterality of the hypothesized effects, we collapsed across hemispheres to create a single ROI. We extracted and averaged parameter estimates across voxels and analyzed them with planned contrasts in a repeated-measures ANOVA context using p<0.05 as a threshold.

##### Studies 2 and 4

In each trial, participants chose one of two decks of cards (see Fig S2). We instructed participants to choose based on their preferences regarding the information they had available for each trial. Participants had two sources of information to rely on. First, each deck was associated with a small monetary payoff (ranging from $0.03 to $0.09 in 2 cents increments). Participants were told that a subset of their choices (5 trials in Study 2, 7 trials in Study 4) would be added to their final compensation for the study. Payoff amounts for each choice varied across trials (and were occasionally equal). Payoff disparities (i.e., the differences in monetary values associated with each deck) followed a quasi-gaussian distribution, such that the most extreme disparities (6 cents in favor of one deck or the other) always appeared the least, twice (Study 2) or thrice (Study 4) for each participant. As disparities grew smaller they gradually became more frequent, with the zero disparity trials (i.e., equivalent monetary values for the two sets) appearing most frequently (5 times in Study 2, 6 times in Study 4). Second, each deck was associated with a specific label (“Typical” versus “Atypical” in Study 2, “Common” versus “Uncommon” in Study 4) that determined the content presented when that deck is selected (stereotypical or counter-stereotypical targets in Study 2, knowledge-consistent or knowledge-violating targets in Study 4). After making their selection, participants saw a face-statement pair corresponding to their choice. For example, if a participant in Study 2 selected the “Typical” deck, they would be presented with a stereotypical target (e.g., in Study 2a a male associated with the statement “CEO of a big company”). Participants then indicated how likely that target was to possess that characteristic on a visual scale of 0 (= “Not at all likely”) to 100 (= “Very likely”; the scale had no other tick marks). The location of the specific labels (right deck versus left deck) was counterbalanced between participants. Study 2a included only faces of men, Study 2b included only faces of women, Study 4a included only the face of Barack Obama and Study 4b included only the face of Donald Trump.

All studies included a demo trial, 4 practice trials, and 25 deck selection trials (32 trials in Study 4). We also included several catch trials to detect if participants were responding without considering the statement presented. The catch trials prompted the participants to slide the bar to the right tick mark or to the left tick mark. Participants had up to 4s to select a card and up to 5s to rate an individual. Participants that did not respond fast enough to more than 20% of the trials got their survey terminated midway through the task. The specific amounts associated with each trial and the face-statement pairs presented per trial were randomized. Following these trials, participants were presented with 4 final manipulation-check trials in which they were asked to select the card with the higher value.

After completing the main task, participants responded to the individual difference scales mentioned above – ASI, SDO, MCP, and NFC. Order of the scales was randomized between participants. No significant correlations between Point of Subjective Equivalence (PSE) and the individual scores on these scales were consistently detected across studies. Finally, participants supplied demographic information (including age, self-reported gender identity, self-reported race with multiple answers enabled and whether they were born in the US). Study 4 also probed participants’ political affiliation (as in Study 3), how much they liked Barack Obama and how much they liked Donald Trump on two separate 0–100 scales. Then, we probed for participants’ intuitions about the goal of the task and fully debriefed them.

### Supplementary Materials Results

#### Behavioral analyses: Neuroimaging studies

Table S1 presents the summaries of participants’ ratings and reaction times in the impression formation task in Study 1. We analyzed rating data with mixed models as implemented in the lme4 package version 1.1-14 (Bates *et al.*, 2014) and the ‘ordinal’ package version 2018.4-19 (Christensen, 2018) for R version 3.4.2 (R Core Team, 2017). As the behavioral data obtained in the impression formation task were ordinal, we analyzed them using cumulative link mixed models (CLMM) with the logit link. To avoid the transformation of raw reaction time data, we used generalized linear models (gLMMs) with the inverse Gaussian identity link (Lo and Andrews, 2015). We included random effects for the intercepts for participants and statements, as well as by-participant random slopes for the fixed effect of stereotypicality. Trials that elicited no response (<1.5% of all trials; no difference between conditions) were excluded from all analyses.

To examine whether our stereotype-consistent and stereotype-violating targets were indeed perceived differently by our participants in Study 1, we tested the effect of our a-priori categorization on behavioral ratings in a CLMM. Stereotypicality was dummy coded with stereotype-neutral trials as the intercept and behavioral ratings were centered on 0. Overall stereotypicality affected ratings, as indicated by leave-one-out model comparison, comparing our model to an intercept only model (*χ^2^*_(2)_ = 44.45, *p* < 0.001). The manipulation worked as anticipated: Stereotype-consistent targets received higher ratings than stereotype-neutral targets (*β*±*SE* = 0.42±0.14, *Z* = 2.99, *p* = 0.003), whereas stereotype-violating targets received lower ratings (*β*±*SE* = −1.22±0.15, *Z* = −8.31, *p* < 0.001).

Stereotypicality did not have a main effect on reaction time for the behavioral ratings (*χ^2^*_(2)_ = 0.35, *p* > 0.8). However, reaction time did vary by the interaction of stereotypicality and behavioral ratings, as indicated by comparing a model with an interaction term to a model without it, *χ^2^*_(2)_ = 133.88, *p* < 0.001); participants were slower to provide responses that were not in line with our a-priori definitions (e.g., indicating that a woman is very likely to be a firefighter or saying that a man is very unlikely to be a CEO); see Table S1 for full descriptive results.

Behavioral results in Study 3 generally replicated the behavioral results we obtained in Study 1 (see Table S3). The outcome of the expectation (whether the target were consistent with or violated the expectation) significantly affected participants’ ratings (*χ^2^*_(1)_ = 73.61, *p* < 0.001). Domain (stereotype-based or person-specific-based knowledge) also affected participants’ ratings (*χ^2^*_(1)_ = 27.58, *p* < 0.001). Interestingly, participants distributed their expectation-based responses differently between the domains (outcome by domain interaction: *χ^2^*_(1)_ = 862.52, *p* < 0.001). Participants used more extreme outcome-congruent ratings for person-specific expectations compared to stereotype-based expectations (see Table S3).

Similar to Study 1, participants’ reaction time did not differ between expectation-consistent or expectation-violating trials (*t =* −0.29, *p =* 0.77), nor between person-specific or stereotype-based expectations (*t* = 1.89, *p* = 0.059). Comparable to Study 1, we observed a significant interaction between outcome and behavioral ratings (*t* = −15.11, *p* < 0.001), an interaction that was qualified by a 3-way interaction with domain (*t* = 6.36, *p* < 0.001). To interpret this interaction, we analyzed responses separately for the two knowledge domains. In both knowledge domains, participants responded faster when their responses were in line with our a-priori definitions (*t* = −9.72 and *t* = −13.81 for stereotype- and person-specific-based expectations, respectively; *p’s* < 0.001).

#### Neuroimaging

In Study 3, in addition to the main findings (see Fig 1 and Table S4), the design allowed us to compare the effects of expectation-consistency with different specific targets, namely, men, women, Trump or Obama. Analyzing activity in the NAcc in a 2 (expectation result) X 2 (content domain) X 2 (specific target) ANOVA yielded, in addition to the main effect of expectation consistency (*F*_(1,29)_ = 21.41, *p* < 0.0001, *η^2^_p_* = 0.42 [0.19–0.58], a main effect of content domain (*F*_(1,29)_ = 4.56, *p* = 0.04, *η^2^_p_* = 0.14 [0.003–0.32]; all other effects p>0.1) such that gender-related trials were associated with increased activity. To complement the findings, we conducted a whole-brain interaction analysis to examine whether any neural regions responded differently to the confirmation or violation of expectations between specific targets. Given the different experimental context between the two content domains, we first examined the interaction of expectations and specific targets within each content domain separately. This analysis yielded 3 regions (p<0.05, FWE-corrected; Table S5 and Figure S3), including the left inferior frontal gyrus (LIFG) and left superior temporal gyrus (LSTG), in which consistency with expectations yielded more activity than their violation only when the statements pertained to Trump (LIFG results: *F*_(1,114.24)_ = 40.12, *p* < 0.0001, *η^2^_p_* = 0.26 [0.15–0.36]) and not to any of the other targets (triple interaction: *F*_(1,29)_ = 33.32, *p* < 0.0001, *η^2^_p_* = 0.53 [0.30–0.66; similar patterns were observed in LSTG]. Disambiguating this triple interaction, we verified the interaction between expectation results and specific targets for leaders: *F*_(1,29)_ = 42.73, *p* < 0.0001, *η^2^_p_* = 0.6 [0.37–0.71]; the parallel interaction within gender content was not significant: *F*_(1,29)_ = 0.55, *p* > 0.4, *η^2^_p_* = 0.02 [0–0.03]). Notably, no main effect of expectation results was observed in these regions (all *p’s* > 0.05). Together, these findings suggest that the consistency with expectations pertaining to Donald Trump evokes an additional process which is not triggered for confirmation of other types of expectations.

#### Behavioral analysis: Online studies

In addition to the analyses reported in the main manuscript (for raw distribution data, see Table S6), in Study 4 we also performed an exploratory pre-registered analysis to examine the relationship between PSEs and support for the specific leader. To that end, we collected from each participant how much they liked Barack Obama and how much they liked Donald Trump on two separate 0–100 scales. Unsurprisingly for online data collection, our participants demonstrated a skewed preference toward Barack Obama (mean Obama liking for participants in Study 4a: 71.89, *S.D*. = 28.68; mean Trump liking for participants in Study 4b: 17.1, *S.D.* = 27.33). We correlated these data with PSEs for the corresponding leader in each study. As can be seen in Figure S4, the data are extremely skewed, with many participants indicating the maximum liking rating for Obama and the minimum liking rating for Trump. As such, we do not provide inferential statistics as these data violate the statistical assumptions. Descriptively, we can see a small positive slope between PSE and liking for each leader. These descriptive trends can be taken to cautiously hint that the more one likes a leader, the more they would be willing to forgo money to see information consistent with the expectation from that leader.

## Supplementary Materials Figures

**Fig S1.**
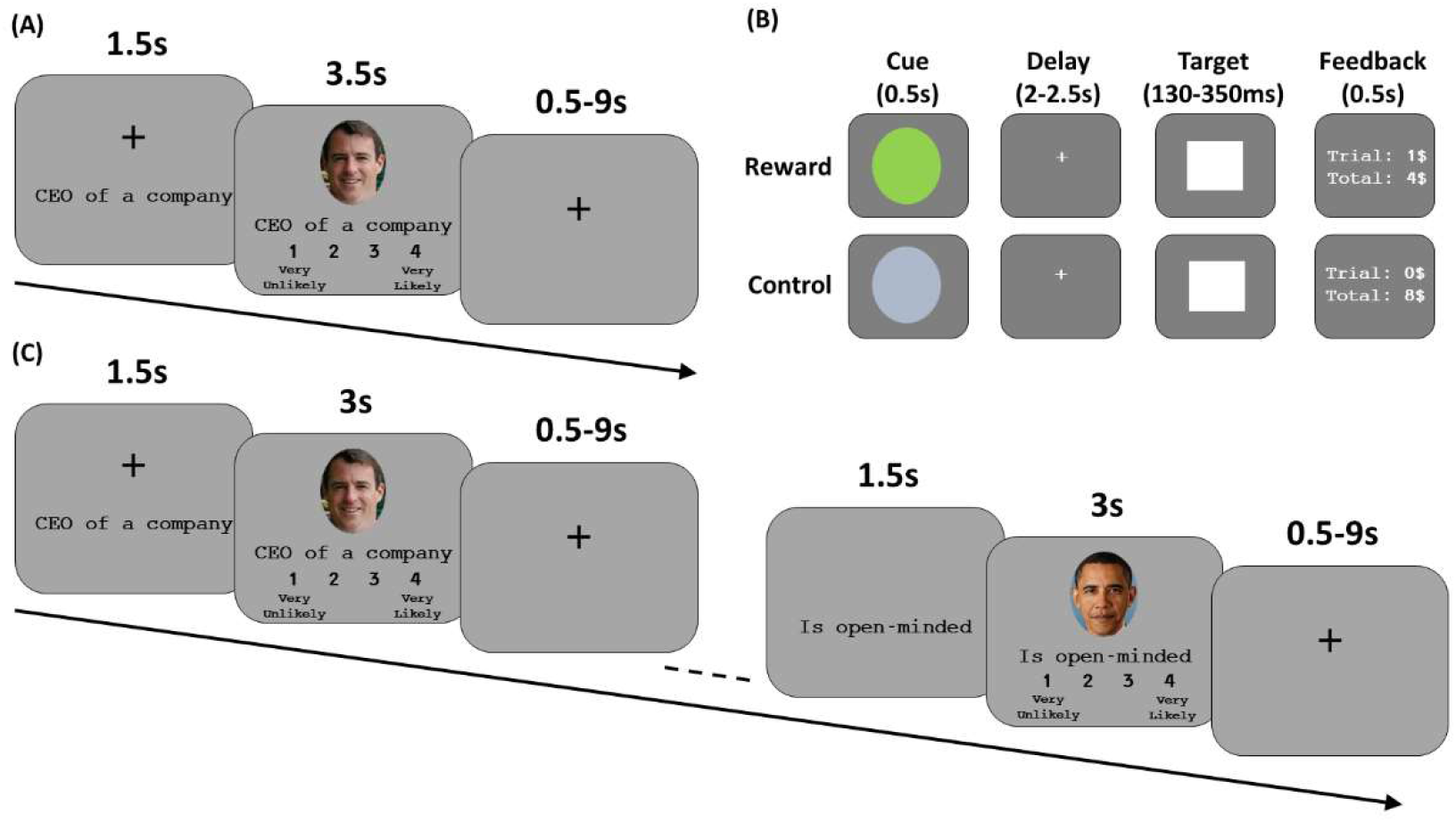
Experimental designs of Studies 1 and 3. (A) In Study 1, participants saw 204 unique trials. Each started with a gender-relevant or irrelevant statement, followed by a target face consistent with, violating or neutral in respect to the displayed statement. Participants indicated how likely the presented person was to have the characteristic described in the statement on a 1 (“very unlikely”) to 4 (“very likely”) scale. (B) Following the impression formation task, participants completed the monetary incentive delay (MID) task. In each trial, participants saw a cue predicting the outcome of a successful response to the target. A green cue always indicated monetary reward, a blue cue always indicated no reward. Participants saw 30 reward cues and 15 no-reward cues. After a randomly jittered delay a target appeared on screen for a brief duration (determined by a 2-up-1-down staircase procedure). Participants received feedback about their performance in each trial and across the entire task. (C) In Study 3, participants rated 240 trials, including 120 stereotype-related targets and 120 person-specific trials. Each content domain (stereotypes versus person-specific) was presented in separate blocks of 15 consecutive trials per type.

**Fig S2.**
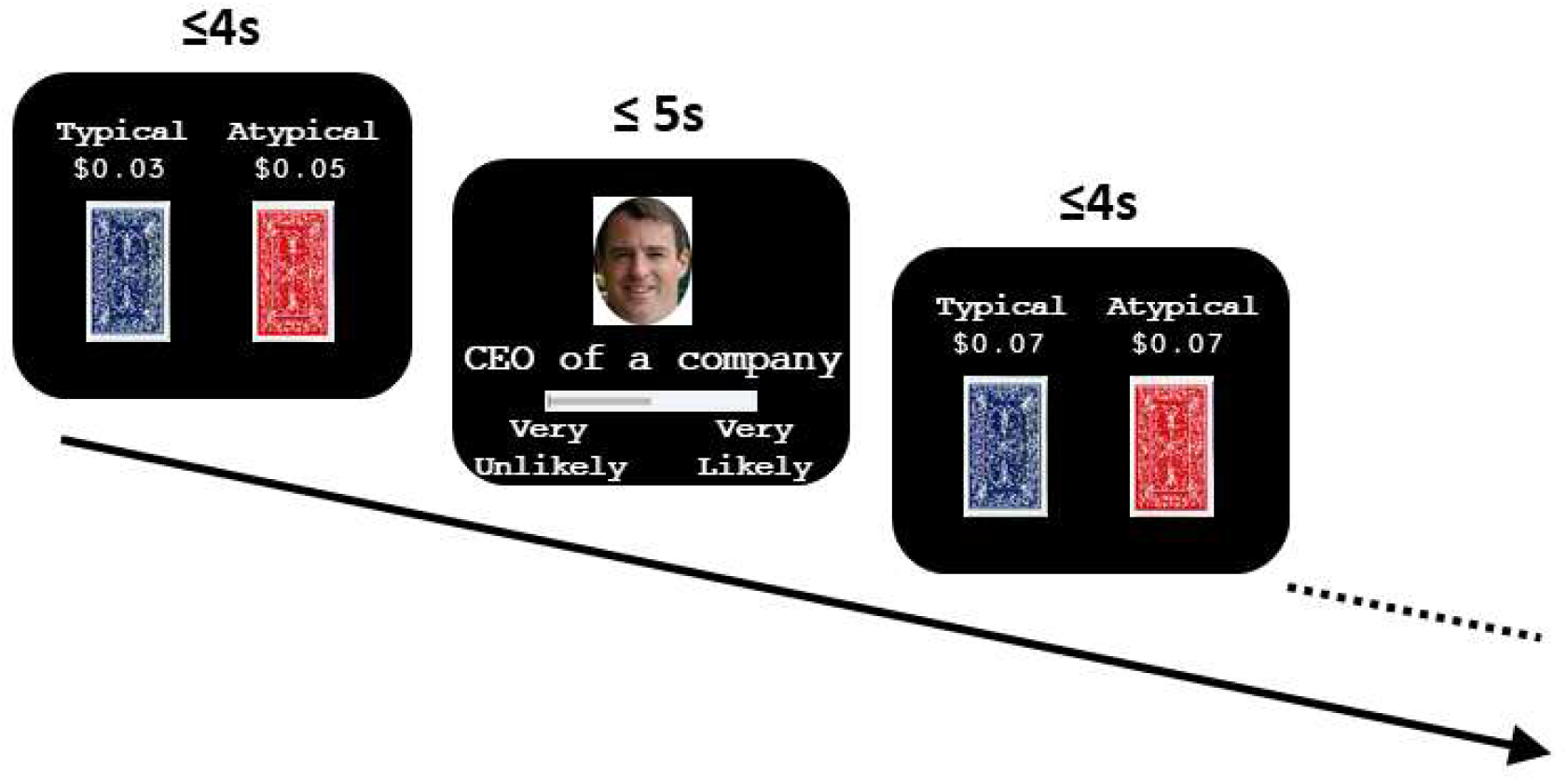
Experimental design of Study 2a. Participants chose which target type to rate (typical, denoting stereotype-confirming targets, versus atypical, denoting stereotype-violating targets). Each target type was associated with a variable amount of money. After deciding which target type to rate, participants then rated a target of the chosen type. The design of studies 2b, 4a and 4b followed this procedure.

**Fig S3.**
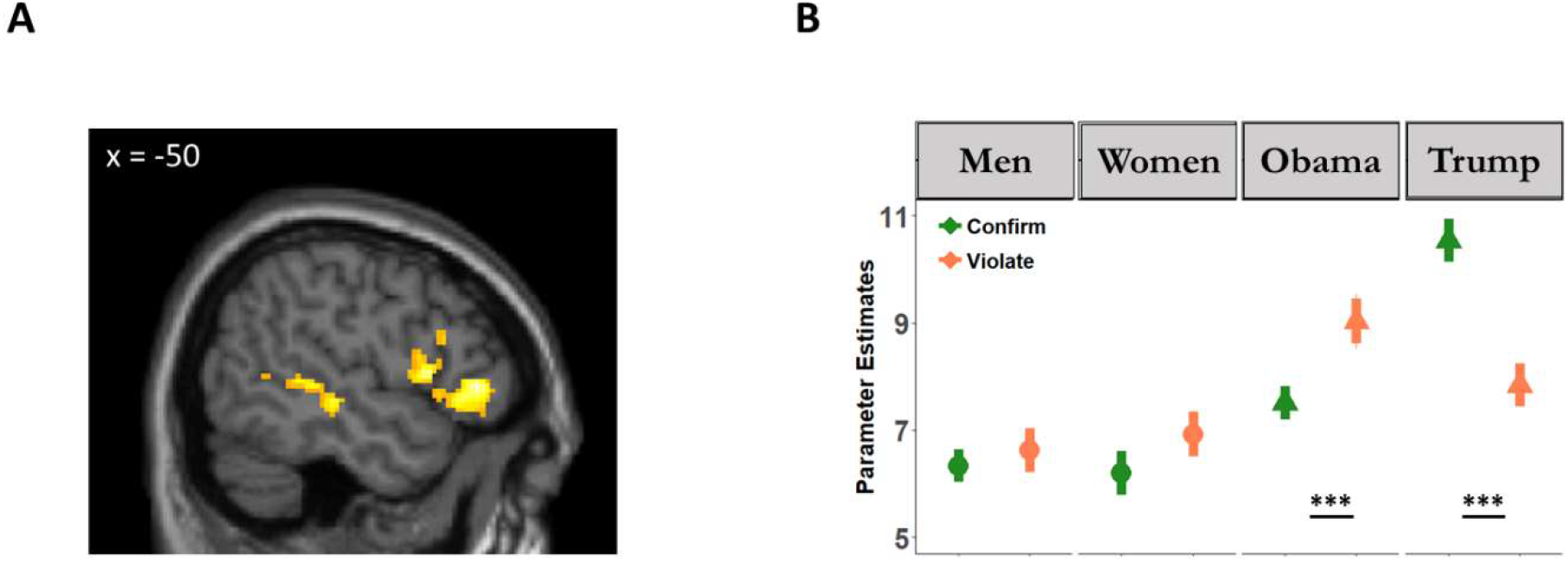
Neural responses as a function of expectancy-outcome and type of target. (A) results of a whole-brain analysis examining the interaction of outcome and type of target (see main text and Table S5 for details). (B) Parameter estimates drawn from the Left Inferior Frontal Gyrus ROI.

**Fig S4.**
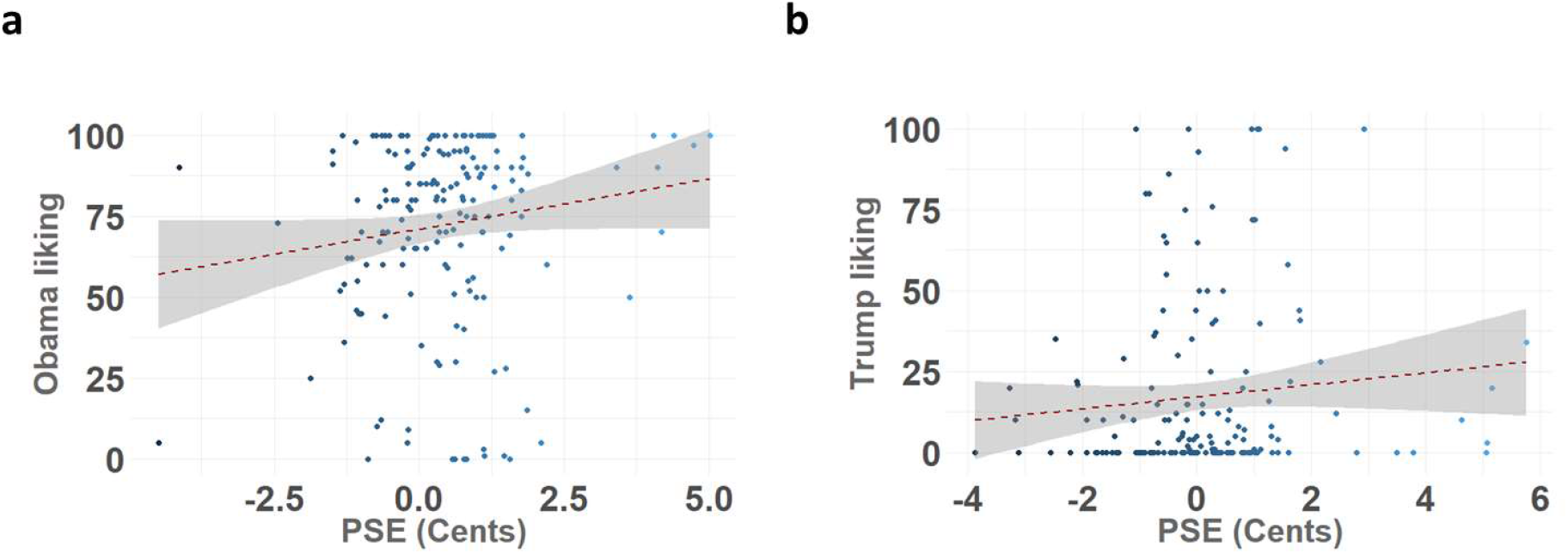
The correlations between the subjective value of expectation-consistent information and the liking of a specific leader in Study 4. The subjective value was calculated as the point of subjective equivalence (PSE) per participant, as detailed in the main text. (a) Results from Study 4a in which participants chose between seeing trials with expectation-consistent and expectation-inconsistent information about Barack Obama. (b) Results from Study 4b in which participants made equivalent choices about trials including Donald Trump. We do not provide inferential statistics as these data are heavily skewed.

**Table S1.** Distribution of participants’ ratings and reaction time (RT, in milliseconds) as a function of stereotypicality in the impression formation task in Study 1. Parentheses indicate standard error of the mean.

|  | *Very Likely* | *Somewhat Likely* | *Somewhat Unlikely* | *Very Unlikely* | *No Response* |
| --- | --- | --- | --- | --- | --- |
| ***Stereotype-confirming (68)*** |  |  |  |  |  |
| Proportion | 0.27 (0.03) | 0.39 (0.03) | 0.23 (0.02) | 0.09 (0.01) | 0.01 (0.005) |
| RT | 1676 (28) | 1845 (27) | 2100 (30) | 1751 (54) |  |
| ***Stereotype-neutral (68)*** |  |  |  |  |  |
| Proportion | 0.18 (0.02) | 0.42 (0.03) | 0.26 (0.02) | 0.12 (0.02) | 0.01 (0.004) |
| RT | 1676 (34) | 1855 (31) | 2047 (35) | 1814 (69) |  |
| ***Stereotype-violating (68)*** |  |  |  |  |  |
| Proportion | 0.07 (0.01) | 0.27 (0.02) | 0.36 (0.02) | 0.29 (0.02) | 0.01 (0.005) |
| RT | 1828 (77) | 1946 (26) | 2015 (31) | 1640 (32) |  |

**Table S2.**
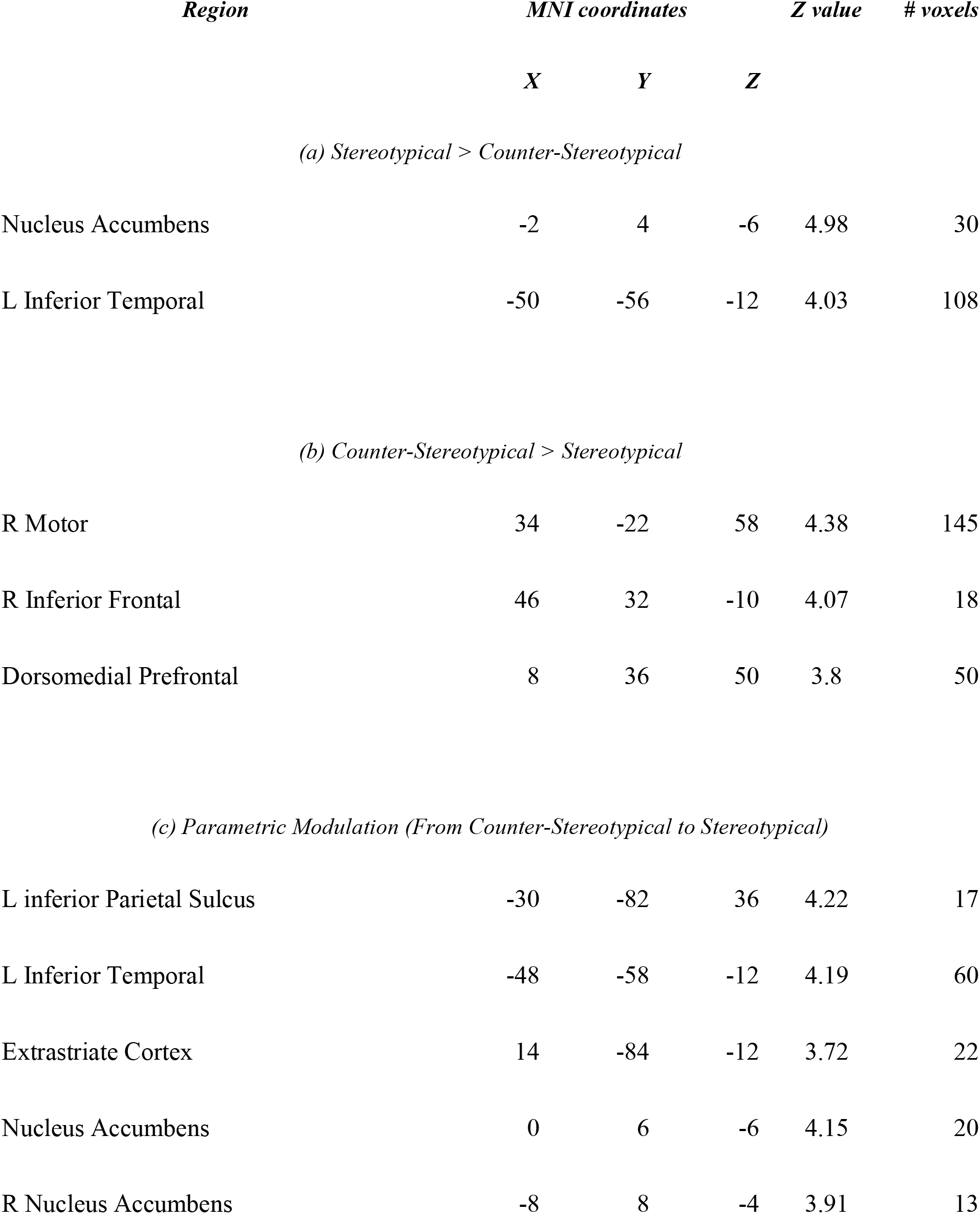
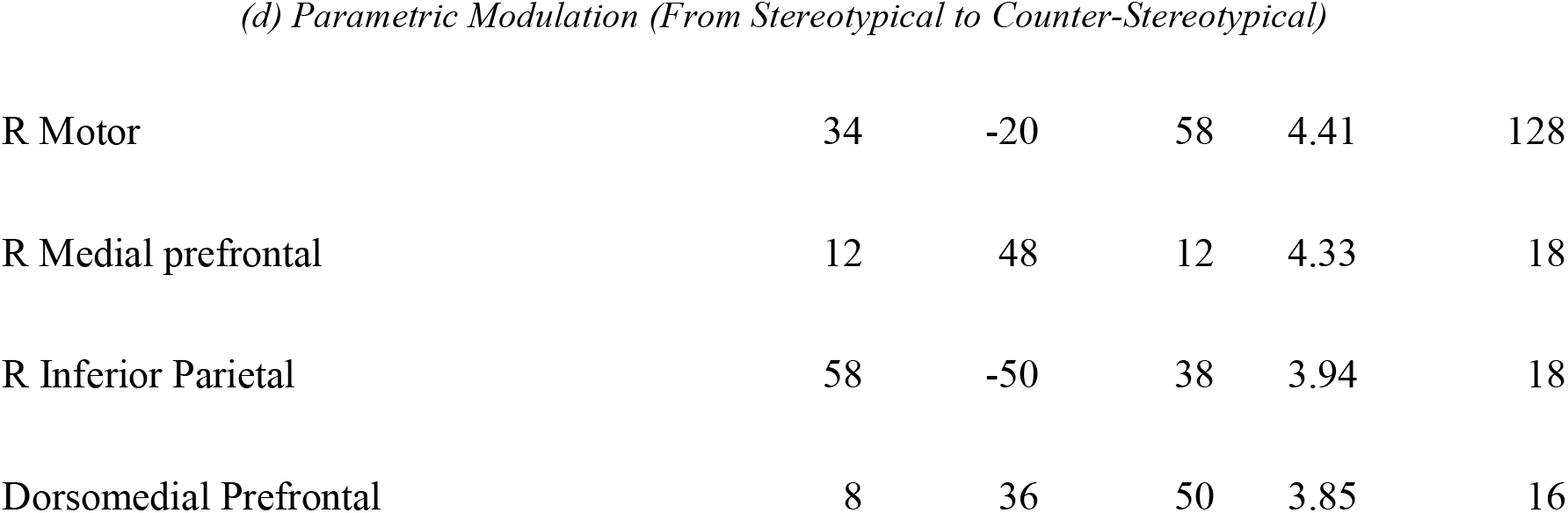
Study 1: Gray matter regions showing differences in activity between stereotypical (stereotype-consistent) and counter-stereotypical (stereotype-violating) targets in model 1 (with no specification of behavioral response; p<0.05, corrected).

| *Region* | *MNI coordinates* |  |  | *Z value* | *# voxels* |
| --- | --- | --- | --- | --- | --- |
|  | *X* | *Y* | *Z* |  |  |
| *(a) Stereotypical > Counter-Stereotypical* |  |  |  |  |  |
| Nucleus Accumbens | -2 | 4 | -6 | 4.98 | 30 |
| L Inferior Temporal | -50 | -56 | -12 | 4.03 | 108 |
| *(b) Counter-Stereotypical > Stereotypical* |  |  |  |  |  |
| R Motor | 34 | -22 | 58 | 4.38 | 145 |
| R Inferior Frontal | 46 | 32 | -10 | 4.07 | 18 |
| Dorsomedial Prefrontal | 8 | 36 | 50 | 3.8 | 50 |
| *(c) Parametric Modulation (From Counter-Stereotypical to Stereotypical)* |  |  |  |  |  |
| L inferior Parietal Sulcus | -30 | -82 | 36 | 4.22 | 17 |
| L Inferior Temporal | -48 | -58 | -12 | 4.19 | 60 |
| Extrastriate Cortex | 14 | -84 | -12 | 3.72 | 22 |
| Nucleus Accumbens | 0 | 6 | -6 | 4.15 | 20 |
| R Nucleus Accumbens | -8 | 8 | -4 | 3.91 | 13 |
| *(d) Parametric Modulation (From Stereotypical to Counter-Stereotypical)* |  |  |  |  |  |
| R Motor | 34 | -20 | 58 | 4.41 | 128 |
| R Medial prefrontal | 12 | 48 | 12 | 4.33 | 18 |
| R Inferior Parietal | 58 | -50 | 38 | 3.94 | 18 |
| Dorsomedial Prefrontal | 8 | 36 | 50 | 3.85 | 16 |

**Table S3.**
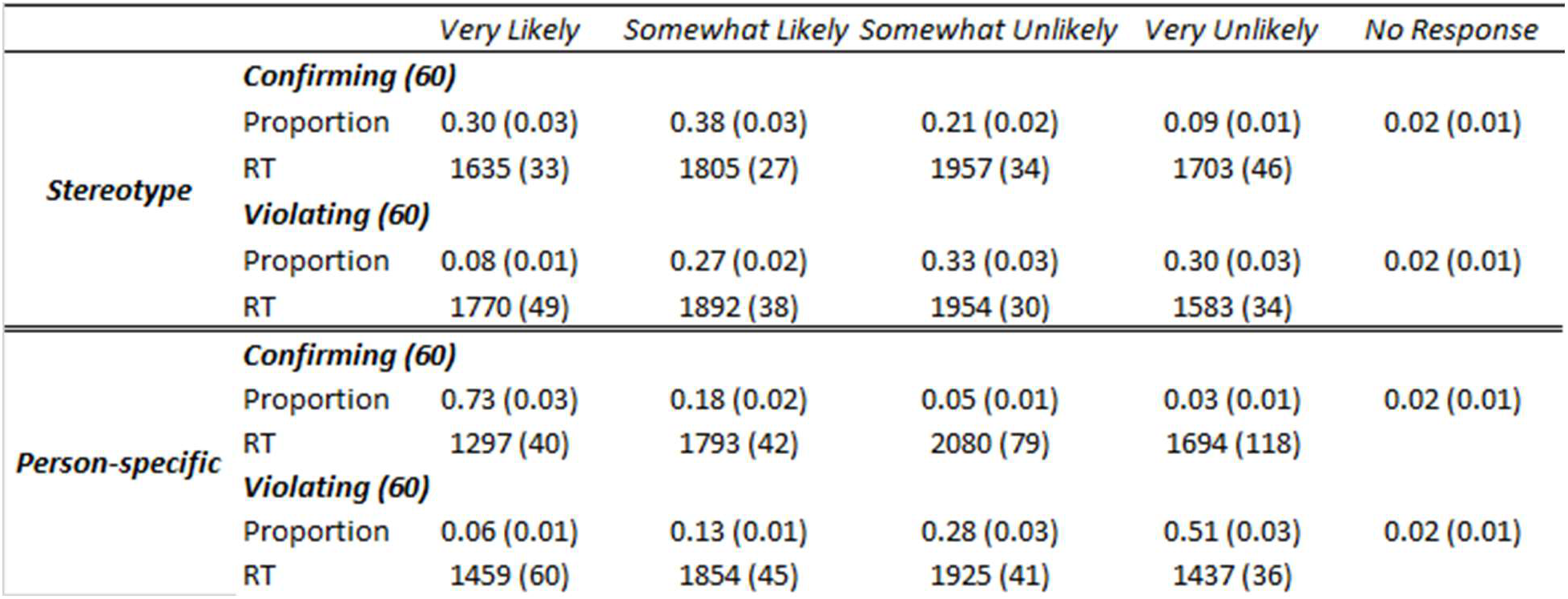
Distribution of participants’ ratings and reaction time (RT, in milliseconds) as a function of expectation domain and expectation-consistency in the impression formation task in Study 3. Parentheses indicate standard error of the mean.

**Table S4.**
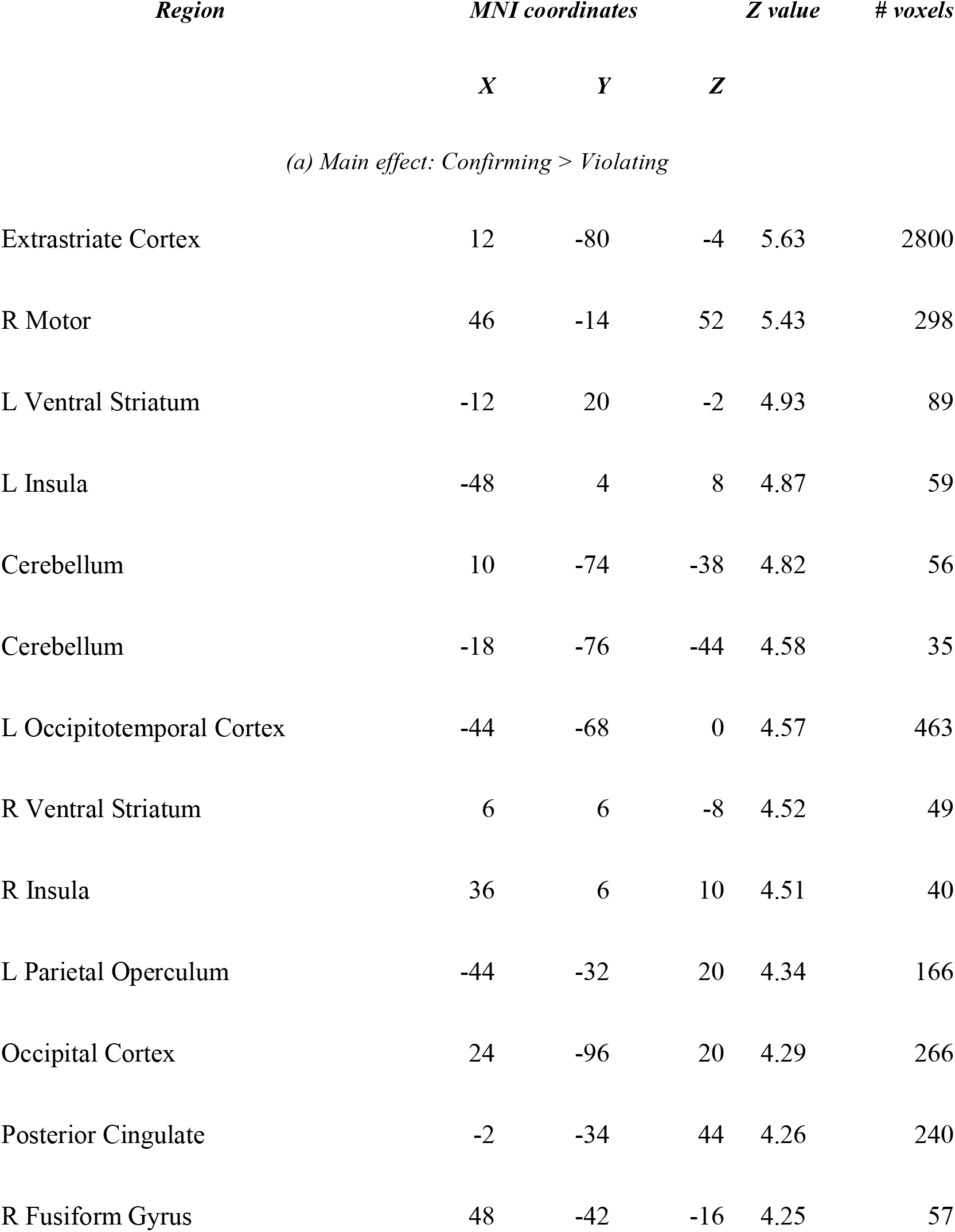
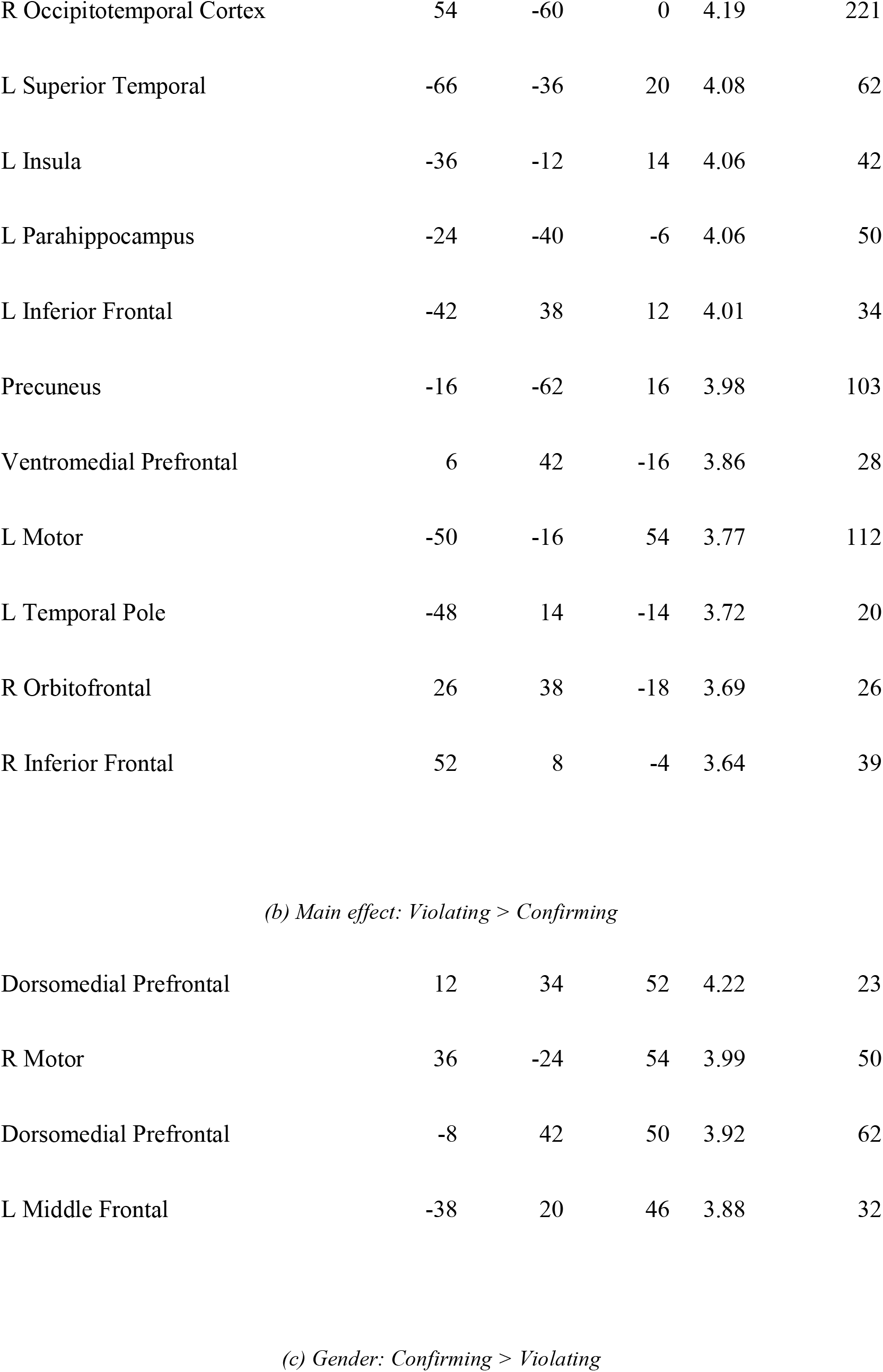
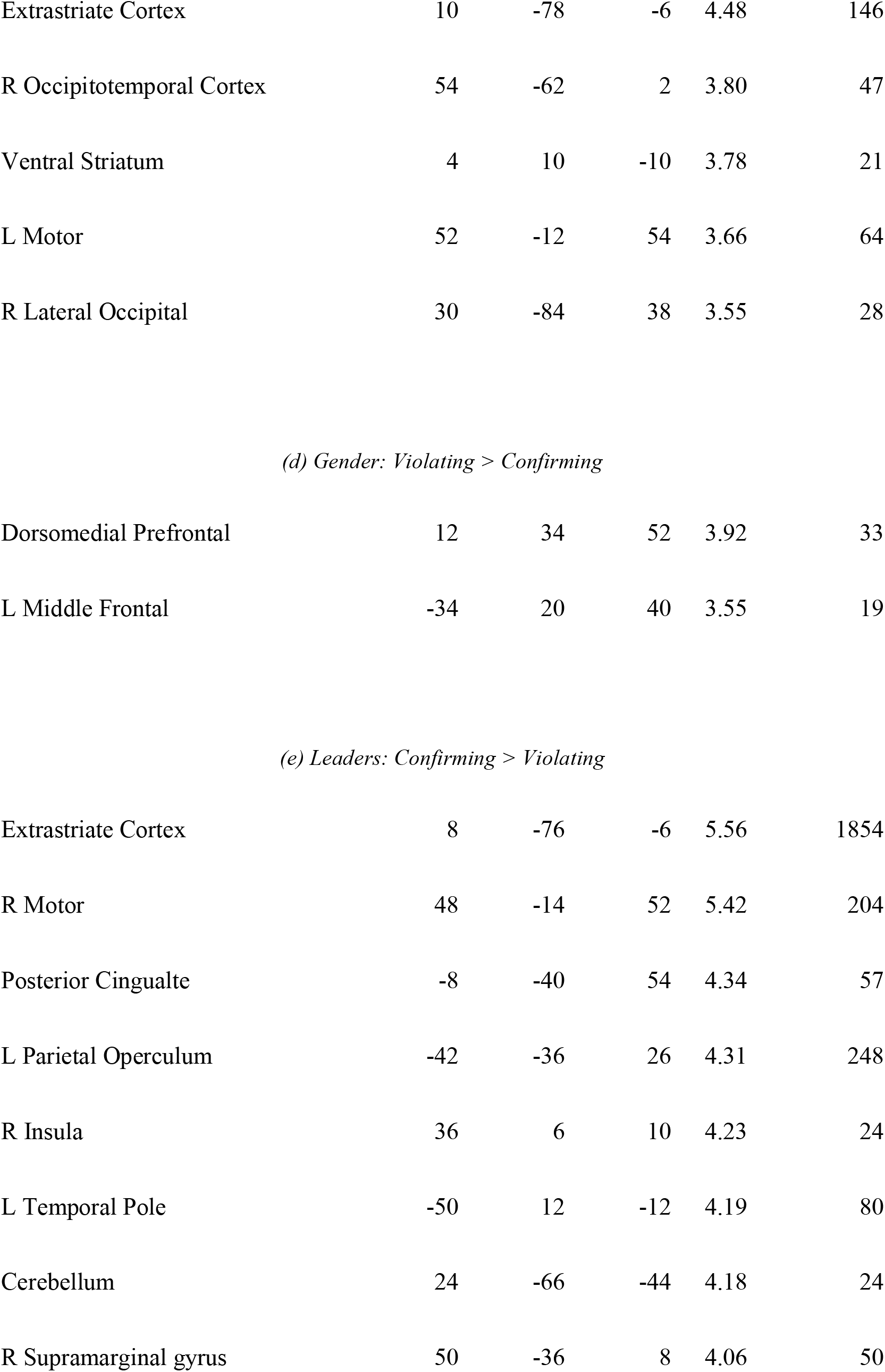
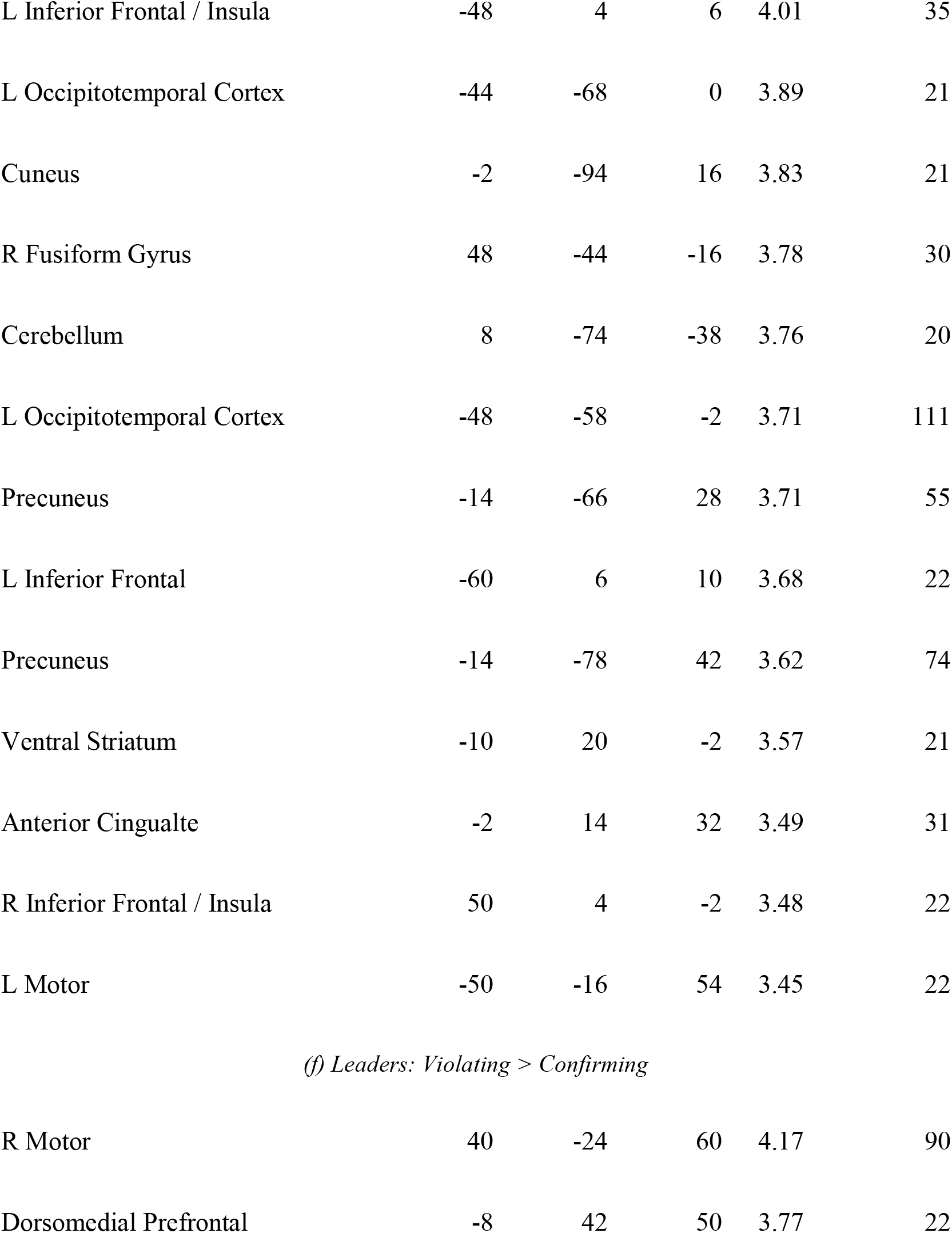
Study 3: Gray matter regions showing differences in activity between expectation-consistent and expectation-violating targets. (a) and (b) portray the results collapsed across content domain, (c) through (f) include the results per content domain (p<0.05, corrected).

**Table S5.**
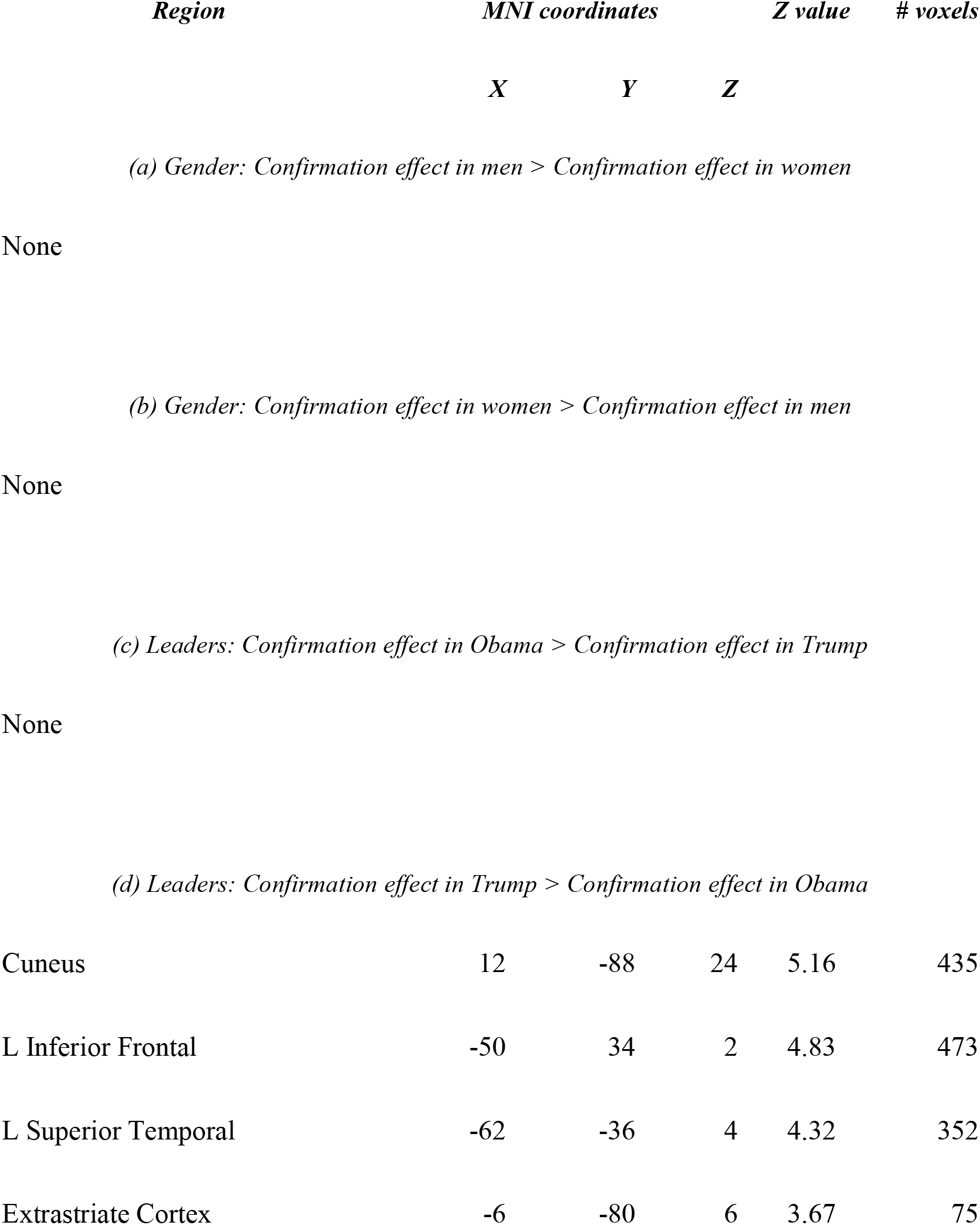
Study 3: Results of a set of two whole-brain analyses examining the interactive effects of expectation-consistency and specific target, done separately for gender and leaders. (p<0.05, corrected).

| *Region* | *MNI coordinates* |  |  | *Z value* | *# voxels* |
| --- | --- | --- | --- | --- | --- |
|  | *X* | *Y* | *Z* |  |  |
| *(a) Gender: Confirmation effect in men > Confirmation effect in women* |  |  |  |  |  |
| None |  |  |  |  |  |
| *(b) Gender: Confirmation effect in women > Confirmation effect in men* |  |  |  |  |  |
| None |  |  |  |  |  |
| *(c) Leaders: Confirmation effect in Obama > Confirmation effect in Trump* |  |  |  |  |  |
| None |  |  |  |  |  |
| *(d) Leaders: Confirmation effect in Trump > Confirmation effect in Obama* |  |  |  |  |  |
| Cuneus | 12 | -88 | 24 | 5.16 | 435 |
| L Inferior Frontal | -50 | 34 | 2 | 4.83 | 473 |
| L Superior Temporal | -62 | -36 | 4 | 4.32 | 352 |
| Extrastriate Cortex | -6 | -80 | 6 | 3.67 | 75 |

**Table S6.**
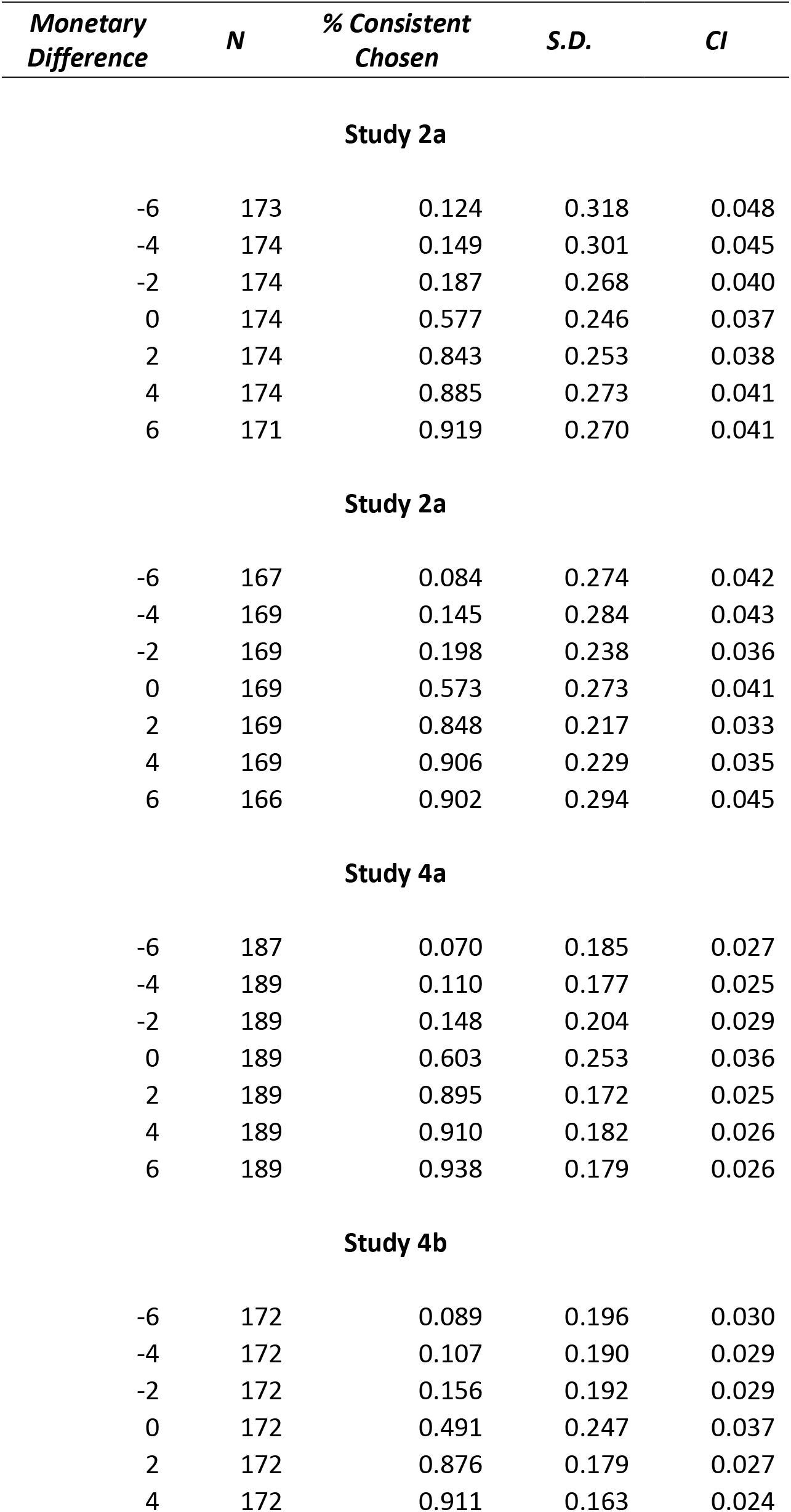
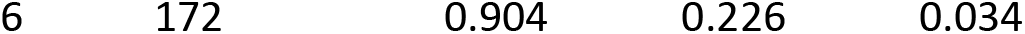
Distribution of proportions of trials in Studies 2 and 4 in which participants chose to see an expectation-confirming target. Data are presented as a function of the difference in monetary value between the two decks of cards.

